# Redefining Interleukin 11 as a regeneration-limiting hepatotoxin

**DOI:** 10.1101/830018

**Authors:** Anissa A. Widjaja, Jinrui Dong, Eleonora Adami, Sivakumar Viswanathan, Benjamin Ng, Brijesh K. Singh, Wei Wen Lim, Jin Zhou, Leroy S. Pakkiri, Shamini G. Shekeran, Jessie Tan, Sze Yun Lim, Mao Wang, Robert Holgate, Arron Hearn, Paul M. Yen, Sonia P. Chothani, Leanne E. Felkin, James W. Dear, Chester L. Drum, Sebastian Schafer, Stuart A. Cook

## Abstract

Acetaminophen (APAP) overdose is a leading cause of untreatable liver failure. In the mouse model of APAP-induced liver injury (AILI), the administration of recombinant human interleukin 11 (rhIL11) is protective. Here we show that the beneficial effect of rhIL11 in the mouse is due to its unexpected and paradoxical inhibition of endogenous mouse IL11 activity. Contrary to the accepted paradigm IL11 is a potent hepatotoxin across species, which is secreted from damaged hepatocytes to drive an autocrine loop of NOX4 and JNK-dependent apoptosis. Mice with hepatocyte-specific *Il11* expression spontaneously develop liver failure whereas those with *Il11ra1* deletion are remarkably protected from AILI. Neutralizing anti-IL11R antibodies administered to moribund mice 10 hours following a lethal APAP overdose results in 90% survival that is associated with very large liver regeneration. Our data overturn a misconception, identify a new disease mechanism and suggest IL11 as a therapeutic target for liver regeneration.

## Main text

Acetaminophen (N-acetyl-p-aminophenol, APAP) is an over-the-counter analgesic that is commonly taken as an overdose (OD) leading to APAP-induced liver injury (AILI), a major cause of acute liver failure^1^. The antioxidant N-acetyl cysteine (NAC) is beneficial for patients presenting early^2^, but there is no drug-based treatment beyond eight hours post-OD and death can ensue if liver transplantation is not possible^3, 4^.

In hepatocytes, APAP is metabolized to N-Acetyl-p-benzochinonimin (NAPQI) which depletes cellular glutathione (GSH) levels and damages mitochondrial proteins leading to reactive oxygen species (ROS) production and JNK activation^5^. ROS-related JNK activation results in a combination of necrotic, apoptotic and other forms of hepatocyte cell death causing liver failure^1, 6, 7^. JNK and ASK1 inhibitors have partial protective effects against AILI in mouse models, but this has not translated to the clinic^8, 9^.

Liver regeneration has fascinated humans since the stories of Prometheus and can be truly profound, as seen after partial hepatic resection in rodents and humans^10, 11^. However, in the setting of AILI, liver regeneration is persistently suppressed resulting in permanent injury and patient mortality. Targeting the pathways that hinder the liver’s extraordinary regenerative capacity may trigger natural regeneration, which could be particularly useful in AILI^12, 13^.

Interleukin 11 (IL11) is a scarcely studied cytokine that is of critical importance for myofibroblast activation and fibrosis of the heart, kidney, lung, and liver^14–16^. It is established that IL11 is secreted from injured hepatocytes and Il11 can be detected at high levels in the serum of the mouse model of AILI, where its expression is considered compensatory and cytoprotective^17^. In keeping with this paradigm, administration of recombinant human IL11 (rhIL11) is effective in treating the mouse model of AILI and also protects against liver ischemia, endotoxemia or inflammation^17–22^. As recently as 2016, rhIL11 has been proposed as a treatment for patients with AILI^23^.

During our studies of liver fibrosis we made the unexpected observation that, in the context of some models of fibro-inflammatory liver disease, IL11 may be detrimental for hepatocyte function^14^. This apparent discrepancy with the previous literature prompted us to look in more detail at the effects of IL11 on hepatocytes independent of fibrosis and we chose to do so in the mouse model of AILI, where Il11 is largely upregulated^17^.

### IL11 drives APAP-induced hepatocyte cell death

As reported previously^17^, we confirmed that AILI was characterized by elevated IL11 serum levels in injured mice (Fig. 1A). We then addressed whether the elevated IL11 serum levels in the mouse AILI model originated in the liver. APAP induced a strong upregulation of hepatic *Il11* transcripts (35-fold, P<0.0001). Bioluminescent imaging of a reporter mouse with *luciferase* cloned into the start codon of *Il11* indicated *IL11* expression throughout the liver (Fig. 1B-C, Extended Data Fig. 1). Western blotting confirmed IL11 upregulation at the protein level across a time course of AILI (Fig. 1D). Experiments using a second reporter mouse with an *EGFP* reporter construct inserted into the 3’UTR of *Il11* (Extended Data Fig. 2) showed that following APAP, IL11 protein is highly expressed in necrotic centrilobular hepatocytes, the pathognomonic feature of AILI, coincident with cleaved caspase 3 (Cl. CASP3) (Fig. 1E).

**Figure 1:**
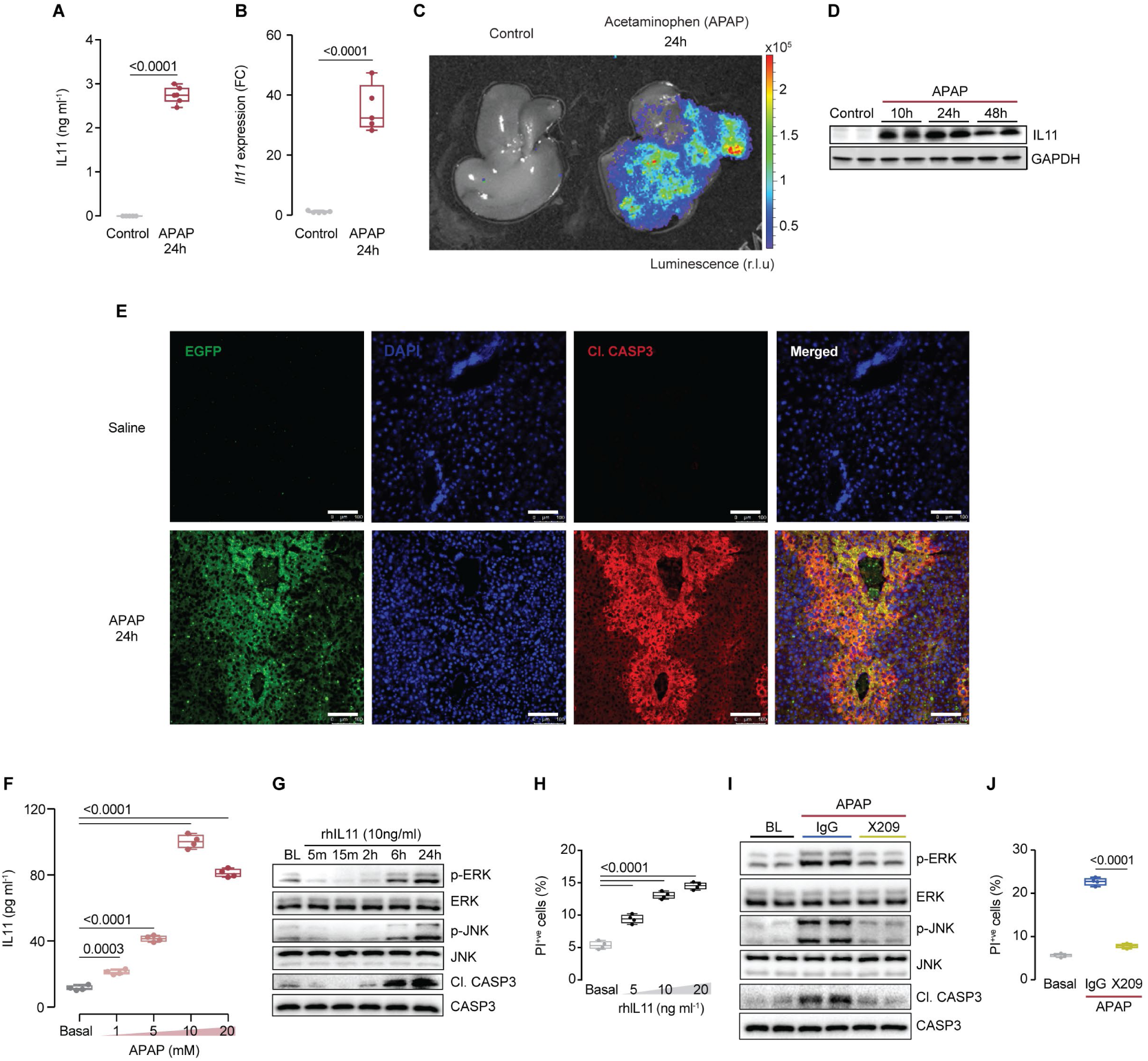
Acetaminophen-induced IL11 secretion from injured hepatocytes causes cell death. (**A**) Serum IL11 levels in APAP-treated mice. (**B**) Liver *Il11* mRNA following APAP injury. (**C**) Representative images of luciferase activity in a liver from control and APAP-challenged *Il11-Luciferase* mice. (**D**) Western blots showing hepatic IL11 expression in APAP-treated mice. (**E**) Representative immunofluorescence images (scale bars, 100 µm) of EGFP and cleaved Caspase3 (Cl. CASP3) expression in the livers of *Il11*-*EGFP* mice post APAP. (**A-E**) APAP, 400 mg kg^-1^. (**F**) ELISA of IL11 secretion from APAP-stimulated hepatocytes. (**G**) Western blots of phosphorylated ERK, JNK and Cl. CASP3 protein and their respective total expression in hepatocytes in response to rhIL11 stimulation. (**H**) Quantification of propidium iodide positive (PI^+ve^) cells from rhIL11-stimulated hepatocytes. (**I**) Western blots showing ERK, JNK, and CASP3 activation status and (**J**) quantification of PI^+ve^ cells in APAP-treated hepatocytes (20 mM) in the presence of IgG or anti-IL11RA (X209; 2 µg ml^-1^). (**F-J**) primary human hepatocytes (**F, H-J**) 24h. (**A, B, F, H-I**) Data are shown as box-and-whisker with median (middle line), 25th–75th percentiles (box), and minimum-maximum values (whiskers). (**A, B**) Two-tailed Student’s *t*-test; (**F, H**) two-tailed Dunnett’s test; (**J**) two-tailed, Tukey-corrected Student’s *t*-test.

Having identified the source of *Il11* upregulation during AILI *in vivo*, we conducted *in vitro* experiments to study underlying mechanisms. Exposure of primary human hepatocytes to APAP resulted in the dose-dependent secretion of IL11 (Fig. 1F). Hepatocytes express interleukin 11 receptor subunit alpha (IL11RA) and it is known that IL11 activates ERK in some cell types^14^, hence we explored the effect of IL11 on ERK and JNK, important in AILI, activation in hepatocytes. IL11 induced late (>6h) and sustained ERK and JNK activation that was concurrent with CASP3 cleavage (Fig. 1G). FACS-based analyses showed dose-dependent IL11-induced hepatocyte cell death (Fig. 1H, Extended Data Fig. 3A). To explore the role of IL11 signaling in APAP-challenged hepatocytes, we used an IL11RA neutralizing antibody (X209)^14^, which inhibited CASP3 cleavage and cell death, as well as ERK and JNK activation (Fig. 1I-J, Extended Data Fig. 3B). While these data confirm the upregulation of IL11 in AILI, they challenge the common perception that this effect is compensatory and protective in the injured liver.

### Species-specific effects of recombinant human IL11

rhIL11 is consistently reported to be protective in rodent models of liver damage^17–20, 23^, yet our studies suggested rhIL11 has the exact opposite effect on human hepatocytes *in vitro* (Fig. 1). This prompted us to test for potential inconsistencies when rhIL11 protein is used in foreign species, as human and mouse IL11 share only 82% protein sequence homology. First, we compared the effects of rhIL11 versus recombinant mouse IL11 (rmIL11) on mouse hepatocytes. While the species-matched rmIL11 stimulated ERK and JNK phosphorylation and induced CASP3 cleavage in mouse hepatocytes, rhIL11 had no effect (Fig. 2A). Similarly, while rmIL11 induced mouse hepatocyte cell death, rhIL11 did not. Indeed, at higher doses rhIL11 trended towards inhibiting mouse hepatocyte death (Fig. 2B). In reciprocal experiments in human hepatocytes, we found that rhIL11 stimulated ERK and JNK signaling and hepatocyte death, whereas rmIL11 did not (Extended Data Fig. 4A-B).

**Figure 2:**
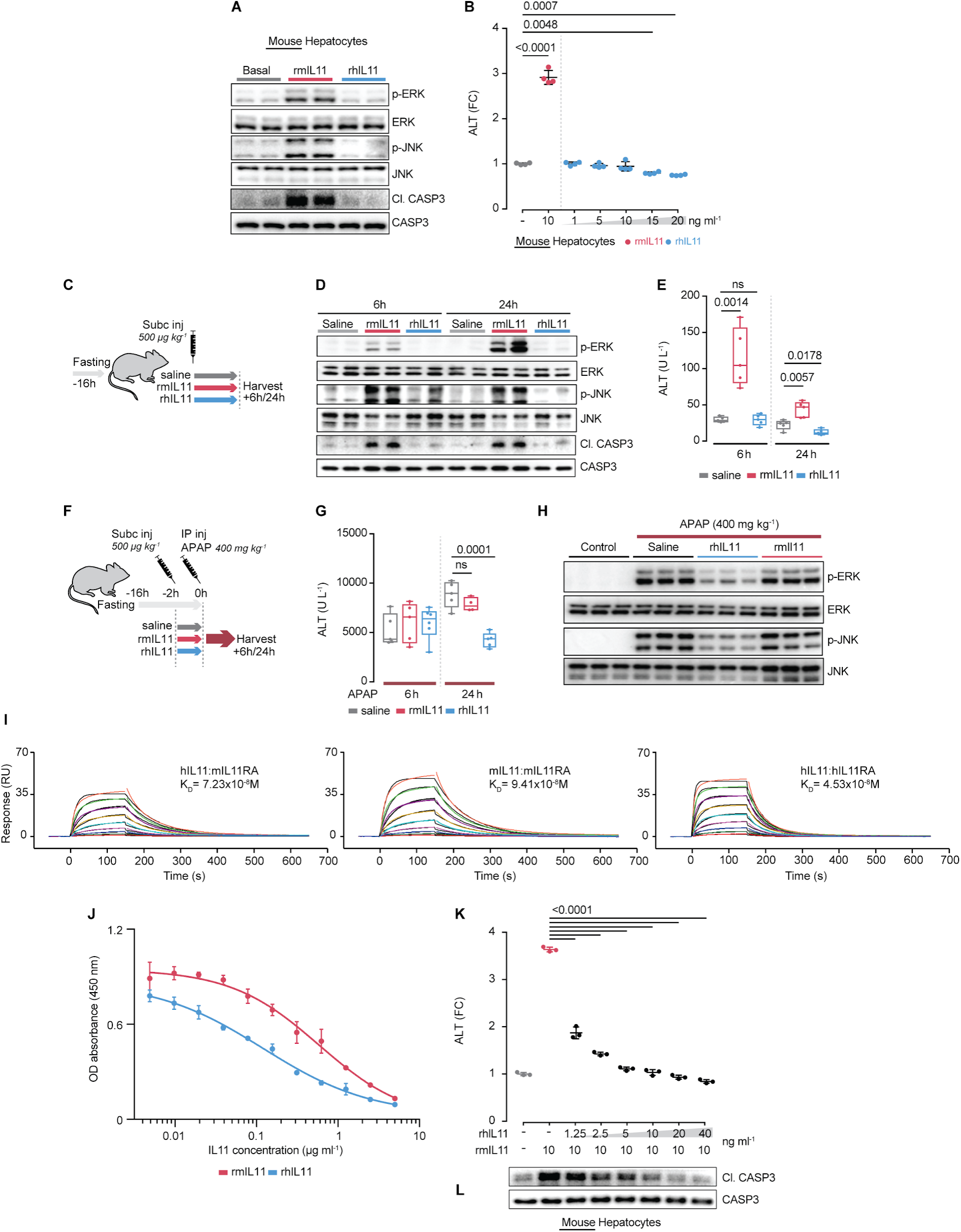
Recombinant human IL11 inhibits mouse IL11 effects in mouse hepatocytes. (**A**) Effect of recombinant human IL11 (rhIL11, 10 ng ml^-1^) or recombinant mouse IL11 (rmIL11, 10 ng ml^-1^) on ERK, JNK and CASP3 activation status in mouse hepatocytes. (**B**) ALT levels in mouse hepatocyte supernatant following stimulation by rmIL11 or by increasing doses of rhIL11. (**C**) Schematic of mice receiving a single subcutaneous injection of either saline, rhIL11, or rmIl11 (500 µg kg^-1^). (**D**) Western blot analysis of hepatic p-ERK, p-JNK, and Cl. CASP3 and (**E**) serum ALT levels of the experiments shown in Fig. 2C. (**F**) Schematic of mice receiving a subcutaneous injection of either saline, rhIL11, or rmIL11 2h prior to APAP OD. Effect of rhIL11 or rmIL11 injection prior to APAP OD on (**G**) serum ALT measurement at 6 and 24h and on (**H**) hepatic ERK and JNK activation at 24h following APAP administration. (**I**) Sensorgrams showing binding of mIL11RA1 to immobilized rhIL11 (left) and rmIL11 (middle), and binding of hIL11RA to rhIL11 (right). The colored lines represent the experimental data; the black lines represent a theoretically fitted curve (1:1 Langmuir). (**J**) Binding of biotinylated rmIL11 to mIL11RA1 in the presence of two-fold dilutions of rmIL11 and rhIL11 by competition ELISA. Dose-dependent inhibition effect of rhIL11 on rmIL11-induced (**K**) ALT secretion and (**L**) CASP3 activation by mouse hepatocytes. (**A, B, K, L)** 24h. (**B, K**) Data are shown as mean±SD; (**E, G**) Data are shown as box-and-whisker with median (middle line), 25th–75th percentiles (box), and minimum-maximum values (whiskers). (**B, K**) Two-tailed, Tukey-corrected Student’s *t*-test; (**E**) two-tailed Student’s *t*-test; (**G**) two-tailed Dunnett’s test. FC: fold change

This showed that the role of IL11 signaling in hepatocyte death is conserved across species, but that recombinant IL11 protein has species-specific effects and does not activate the pathway in foreign species. We tested this hypothesis *in vivo* by injecting either rmIL11 or rhIL11 into mice (Fig. 2C). Injection of rmIL11 resulted in gradual ERK and immediate JNK activation. In contrast, rhIL11 had no effect on ERK or JNK phosphorylation (Fig. 2D). Injection of rmIL11 also caused liver damage with elevated ALT and AST (Fig. 2E, Extended Data Fig. 4C). In stark contrast, rhIL11 injection in naive mice was associated with slightly lower ALT and AST levels 24h post-injection (ALT, P=0.018; AST, P=0.0017).

To follow up on the potential protective effect of rhIL11 in the mouse, we performed a protocol similar to the AILI study of 2001^20^, where rhIL11 was injected into the mouse after APAP OD (Fig. 2F). This confirmed that rhIL11 reduces the severity of AILI in mice (reduction: ALT, 52%, P=0.0001; AST, 39%, P<0.0001), whereas species-matched rmIL11 was not protective in the mouse (Fig. 2G, Extended Data Fig. 4D). The therapeutic effect of rhIL11 was accompanied by a reduction in hepatic ERK and JNK activation (Fig. 2H), which shows that rhIL11 blocks IL11-driven signaling pathways in the liver similar to IL11RA antibodies (Fig. 1I).

Using surface plasmon resonance (SPR), we found that rhIL11 binds to mouse interleukin 11 receptor alpha chain 1 (mIL11RA1) with a K_D_ of 72 nM, which is slightly stronger than the rmIL11:mIL11RA1 interaction (94 nM) and close to that reported previously for rhIL11:hIL11RA (50 nM), which we reconfirmed (Fig. 2I, Extended Data Fig. 4E)^24^. We then performed a competition ELISA assay and found that rhIL11 competed with rmIL11 for binding to mIL11RA1 and was a very effective blocker as suggested by the higher affinity to mIL11RA1 (Fig. 2J). In mouse hepatocytes, rhIL11 was a potent, dose-dependent inhibitor of rmIL11-induced signaling pathways and cytotoxic activity (Fig. 2K-L, Extended Data Fig. 4F). Thus, paradoxically, foreign rhIL11 acts as a neutralizer of mouse IL11 both *in vitro* as *in vivo* and these observations challenge our understanding of the role of IL11 in liver injury and in disease more broadly.

### Hepatocyte-specific expression of *Il11* causes spontaneous liver failure

To test the effects of endogenous mouse IL11 secreted from hepatocytes *in vivo*, we expressed an *Il11* transgene specifically in hepatocytes by injecting *Rosa26^Il11/+^* mice^15, 16^ with AAV8 virus encoding an albumin promoter-driven Cre construct (*Il11*-Tg mice, Fig. 3A). Three weeks after transgene induction, *Il11*-Tg mice had grossly abnormal and smaller (38%, P<0.0001) livers with elevated serum ALT and AST levels, while other organs were unaffected (Fig. 3B-D, Extended Data Fig. 5 A-B). Histologically, there was marked portal vein dilatation and blood accumulation in the sinusoids - suggestive of a sinusoidal obstruction syndrome - as well as infiltrates around the portal triad (Fig. 3E, Extended Data Fig. 5C). Molecular analyses of *Il11*-Tg livers revealed activation of ERK, JNK, and CASP3 cleavage along with increased pro-inflammatory gene expression (Fig. 3F, Extended Data Fig. 5D-E). Thus secretion of IL11 from hepatocytes, as seen with APAP toxicity (Fig. 1), is hepatotoxic.

**Figure 3:**
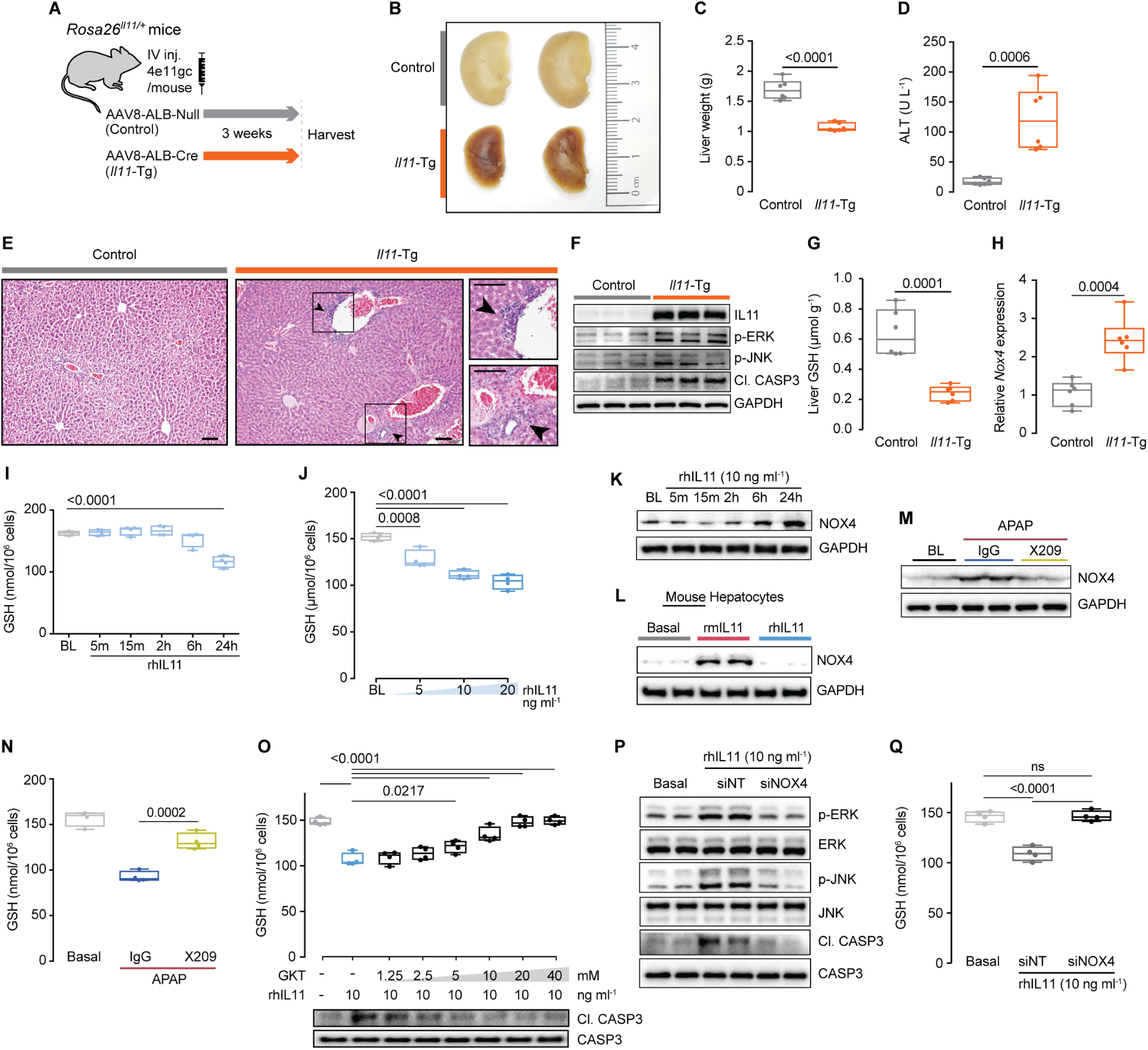
IL11 causes liver failure through NOX4-dependent glutathione depletion. (**A**) Schematic of *Rosa26^Il11/+^* mice receiving a single intravenous injection of either AAV8-ALB-Null (control) or AAV8-ALB-Cre (*Il11*-Tg) to specifically induce *Il11* overexpression in albumin-expressing cells (hepatocytes); ALB: ALBUMIN. (**B**) Representative gross anatomy of livers, (**C**) liver weights, (**D**) serum ALT levels, (**E**) representative H&E-stained liver images (scale bars, 100 µm), (**F**) western blotting of p-ERK, p-JNK, and Cl. CASP3, (**G**) liver GSH levels, and (**H**) *Nox4* mRNA expression levels in control and *Il11*-Tg mice 3 weeks after injection. (**I**) Time course GSH levels, (**J**) dose-dependent decrease in GSH levels, and (**K**) western blots showing increased NOX4 protein expression in rhIL11-treated primary human hepatocytes. (**L**) Western blots of NOX4 in rhIL11 or rmIL11-stimulated mouse hepatocytes. (**M**) Western blots of NOX4 expression and (**N**) GSH levels in IgG and X209-treated APAP-stimulated human hepatocytes (20 mM). (**O**) Dose-dependent inhibition effect of GKT-13781 on GSH levels and CASP3 activation in rhIL11-stimulated human hepatocytes. Effect of siNOX4 on rhIL11-induced (**P**) ERK, JNK, and CASP3 activation and (**Q**) GSH depletion levels in human hepatocytes. (**I-Q**) rhIL11/rmIL11 (10 ng ml^-1^, unless otherwise specified), APAP (20 mM), IgG/X209 (2 µg ml^-1^), siNT (non-targeting siRNA control)/siNOX4 (50 nM). (**I-K, M-Q**) primary human hepatocytes, (**L**) primary mouse hepatocytes. (**J, L-Q**) 24h. (**C-D, G-J, N, O, Q**) Data are shown as box-and-whisker with median (middle line), 25th–75th percentiles (box), and minimum-maximum values (whiskers). (**C-D, G-H**) Two-tailed Student’s *t*-test; (**I-J**) two-tailed Dunnett’s test; (**N**, **O, Q**) two-tailed, Tukey-corrected Student’s *t*-test.

### IL11 stimulates NOX4-mediated reactive oxygen species production

IL11 signaling is required for APAP-driven JNK activation *in vitro* (Fig. 1I-J), which is known to follow ROS production and GSH depletion. We examined liver GSH levels in *Il11*-Tg mice and found they were diminished (62%, P<0.0001), indicating that IL11 signaling - directly or indirectly - induces ROS (Fig. 3G).

In fibroblasts, the expression of *NOX4*, an NADPH oxidase, and source of ROS, is strongly associated with *IL11* expression^15, 25^, and hepatocyte-specific *Nox4* deletion prevents pathological activation of JNK^26^. Therefore, we investigated the relationship between IL11, NOX4, and ROS in greater detail. In *Il11*-Tg mice, hepatic *Nox4* expression was upregulated (Fig 3H). In primary human hepatocytes, IL11 stimulated dose-dependent GSH depletion over a time course that mirrored ERK and JNK activation and was accompanied by NOX4 upregulation (Fig. 1G, Fig. 3I-K). As expected, only species-specific IL11 induced NOX4 upregulation and lowered GSH levels (Fig. 3L, Extended Data Fig. 6A-D).

APAP stimulation also resulted in NOX4 upregulation in hepatocytes, coincident with depletion in hepatocyte GSH levels, which was blocked with the anti-IL11RA antibody X209 (Fig. 3 M-N). We reconsidered the effect of rhIL11 in inhibiting endogenous IL11-induced cell death in mouse hepatocytes (Fig. 2J-K) and found clear, dose-dependent effects of rhIL11 in restoring GSH levels in rmIL11 stimulated mouse cells (Extended Data Fig. 7A). Similarly, rhIL11 restored APAP-induced GSH depletion in the mice, while rmIL11 did not (Extended Data Fig. 7B). GKT-13781, a specific NOX4 inhibitor, prevented IL11-stimulated GSH depletion, CASP3 activation and cell death in a dose-dependent manner (Fig. 3O, Extended Data Fig. 8A-B). The specificity of pharmacological inhibition of NOX4 was confirmed using siRNA, which prevented IL11-induced hepatotoxicity (Fig. 3P-Q, Extended Data Fig. 9A-B). Together these data show that IL11-stimulated NOX4 activity, which could also impact mitochondrial ROS, is important for GSH depletion in the context of AILI.

### Hepatocyte-specific deletion of *Il11ra1* prevents APAP-induced liver failure

To delete *Il11ra1* specifically in adult mouse hepatocytes we created *Il11ra1* conditional knockouts (CKOs) by injecting AAV8-ALB-Cre virus to mice homozygous for LoxP-flanked *Il11ra1* alleles, along with wildtype controls. Three weeks after viral infection, control mice and CKOs were administered APAP (400 mg kg^-1^) (Fig. 4A). The day after APAP administration, gross anatomy revealed small and discolored livers in control mice, whereas livers from CKO mice looked normal (Fig. 4B). Histology showed typical and extensive centrilobular necrosis in control mice, which was not observed in CKOs (Fig. 4C).

**Figure 4:**
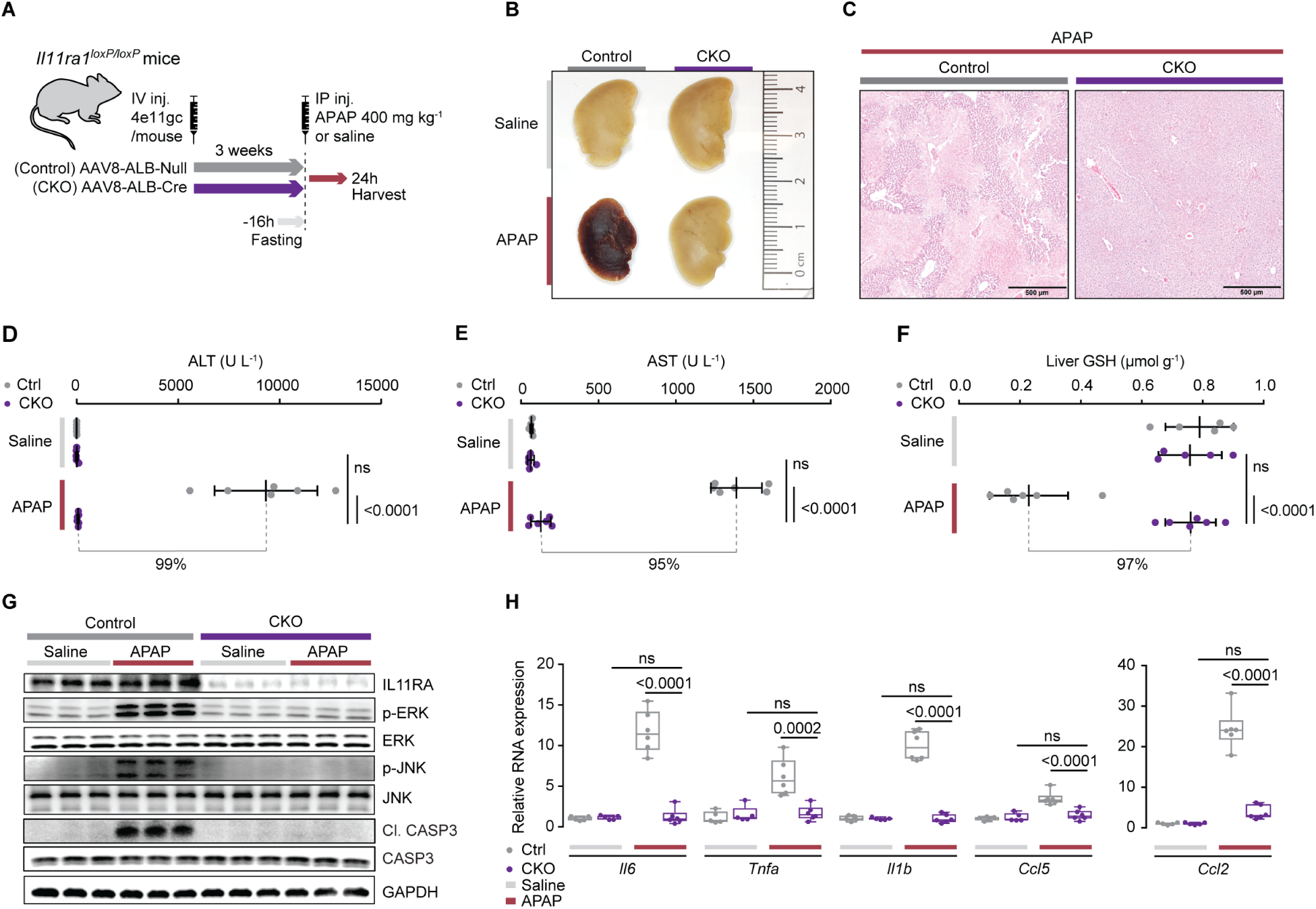
Hepatocyte-specific *Il11ra1* deletion protects mice from APAP-induced liver damage. (**A**) Schematic of induction of APAP injury in *Il11ra1^loxP/loxP^* mice. *Il11ra1^loxP/loxP^* mice were intravenously injected with either AAV8-ALB-Null (control) or AAV8-ALB-Cre (CKO) to specifically delete *Il11ra1* in hepatocytes. Overnight-fasted control and CKO mice were injected with APAP (400 mg kg^-1^) or saline, 3 weeks following virus administration. ALB: Albumin. (**B**) Representative liver gross anatomy and (**C**) H&E images (scale bars, 500 µm) from saline and APAP-injected control and CKO mice. (**D**) Serum ALT levels, (**E**) serum AST levels, (**F**) liver GSH levels, (**G**) western blots of IL11RA, p-ERK, ERK, p-JNK, JNK, Cl. CASP3, CASP3 and GAPDH, and (**H**) relative liver mRNA expression levels of proinflammatory genes. (**D-F, H**) Data are shown as box-and-whisker with median (middle line), 25th–75th percentiles (box), and minimum-maximum values (whiskers); Sidak-corrected Student’s *t*-test.

It was striking that CKO mice had 99% and 95% lower ALT and AST levels, respectively, as compared to controls and GSH levels that were similar to baseline. Both groups had similar levels of APAP and APAP-Glutathione (APAP metabolite) in the serum and thus *Il11ra1* deletion does not impact APAP metabolism (Fig. 4D-F, Extended Data Fig. 10A-B). ERK and JNK activation was observed in control mice, but not in the CKOs (Fig. 4G). Deletion of the receptor in hepatocytes also significantly reduced inflammatory markers, suggesting that inflammation in AILI is secondary to parenchymal injury. (Fig 4H). Taken together, these data show a dominant role for hepatocyte-specific IL11 signaling in the pathogenesis of AILI. The fact that *Il11ra1* deletion in hepatocytes is sufficient to protect from APAP OD indicates that free soluble Il11RA1 in the serum or receptor shedding from other cellular sources does not contribute to disease pathogenesis via *trans*-signaling.

### Effects of anti-IL11RA administration early during APAP-induced liver injury

We next tested if therapeutic inhibition of IL11 signaling was effective in mitigating AILI by administering anti-Il11RA (X209) antibody^14^. Initially, we performed a preventive treatment by injecting X209 or control antibody (10 mg kg^-1^) 16h prior to APAP. This approach reduced serum markers of liver damage by over 70%, largely restored hepatic GSH levels, and limited histological evidence of centrilobular necrosis (Fig. 5A-D, Extended Data Fig. 11A).

**Figure 5:**
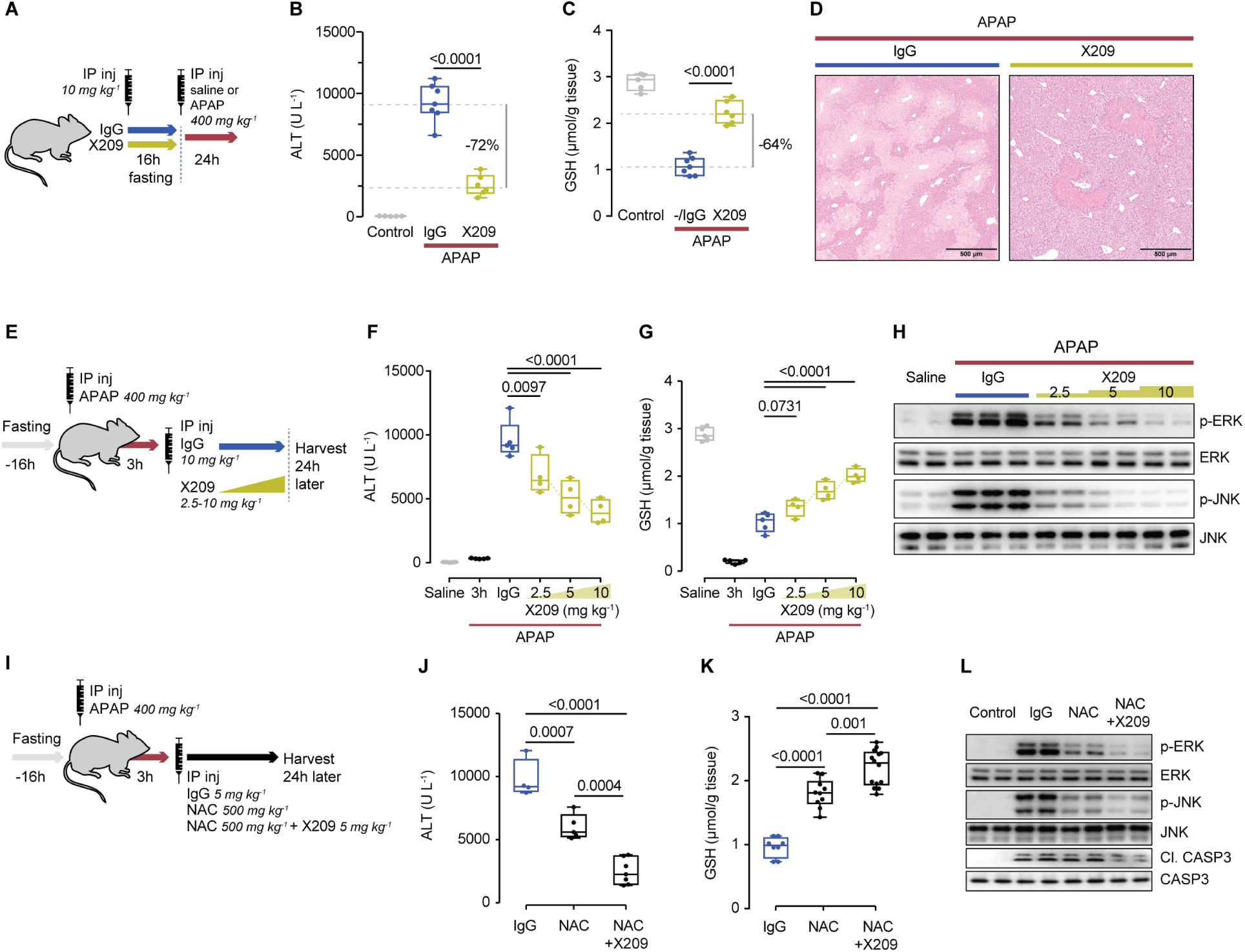
Treatment of APAP-induced liver damage with anti-IL11RA antibody and/or NAC. (**A**) Schematic of anti-IL11RA (X209) preventive dosing in APAP OD mice; X209 or IgG (10 mg kg^-1^) was administered at the beginning of fasting period, 16h prior to APAP (400 mg kg^-1^) injection; control mice received saline injection. (**B**) Serum ALT levels, (**C**) representative H&E images (scale bars, 500 µm), and hepatic GSH levels for the experiments shown in Fig. 5A. (**E**) Schematic of anti-IL11RA (X209) dose finding experiments; X209 (2.5-10 mg kg^-1^) or IgG (10mg kg-1) was administered to mice 3h following APAP injection. (**F**) Serum ALT levels (the values of saline are the same as those used in **5B**), (**G**) hepatic GSH levels, and (**H**) Western blots of hepatic ERK and JNK activation from experiments shown in Fig. 5E. (**I**) Schematic showing therapeutic comparison of X209 and N-acetyl-cysteine (NAC, 500 mg kg ^-1^) alone or in combination with X209 (5 mg kg^-1^). Overnight-fasted mice were treated with IgG, NAC, or NAC+X209 3h post APAP injection for data shown in (**J-L**). Effect of NAC, NAC+X209 treatment on (**H**) serum ALT levels, on (**I**) hepatic GSH levels, and on (**J**) p-ERK, p-JNK, and Cl. CASP3 expression levels (**B, C, F, G, J, K**) Data are shown as box-and-whisker with median (middle line), 25th–75th percentiles (box), and minimum-maximum values (whiskers); two-tailed, Tukey-corrected Student’s *t*-test.

Next, we administered anti-IL11RA therapy in a therapeutically-relevant mode by giving antibody 3h after APAP, a time point by which APAP metabolism and toxicity is established and after which most interventions have no effect in the mouse model of AILI (Fig. 5E)^9^. X209, across a range of doses (2.5-10 mg kg^-1^), inhibited AILI with dose-dependent improvements in markers of liver damage and in hepatic GSH levels. Reduced JNK and ERK activation confirmed dose-dependent target coverage (Fig. 5F-H, Extended Data Fig. 11B).

Lastly, we determined whether inhibiting IL11 signaling had added value when given in combination with the current standard of care, NAC, 3h after APAP dosing (Fig. 5I). Administration of NAC alone reduced serum levels of ALT and AST. However, NAC in combination with X209 was even more effective than either NAC or X209 alone (ALT reduction: NAC, 38%, P=0.0007; X209, 47%, P<0.0001; NAC+X209, 75%; P<0.0001) (Fig. 5F, J, Extended Data Fig. 11C). At the molecular level, the degree of ERK and JNK inhibition with NAC or NAC together with X209 mirrored the magnitude of ALT reduction in the serum and the restoration of hepatic GSH levels (Fig. 5K-L). As such, anti-IL11RA therapy has added benefits when given in combination with the current standard of care.

### Liver regeneration with anti-IL11RA therapy

For patients presenting to the emergency room 8h or later after APAP OD there is no effective treatment. This prompted us to test anti-IL11RA 10h after APAP (400 mg kg^-1^) administration to mice (Fig. 6A). Given the accelerated metabolism of APAP in the mouse, therapy at 10h in this model is equivalent to the treatment of a human up to 24h post-APAP OD. We quantified APAP and APAP-Glutathione in serum by mass spectrometry and found levels to be elevated compared to saline-treated controls and equivalent between experimental groups, as expected (Extended Data Fig. 12A-B). Analysis of gross anatomy, histology and serum IL11, ALT and AST levels revealed that X209 largely reversed liver damage by the second day after APAP, whereas IgG treated mice had profound and sustained liver injury (Fig. 6B-E, Extended Data Fig. 13A). The therapeutic antibody effectively blocked ERK and JNK activation throughout the course of the experiment and this preceded a reduction in cleaved CASP3 at 24h (Fig. 6F, Extended Data Fig. 13B).

**Figure 6:**
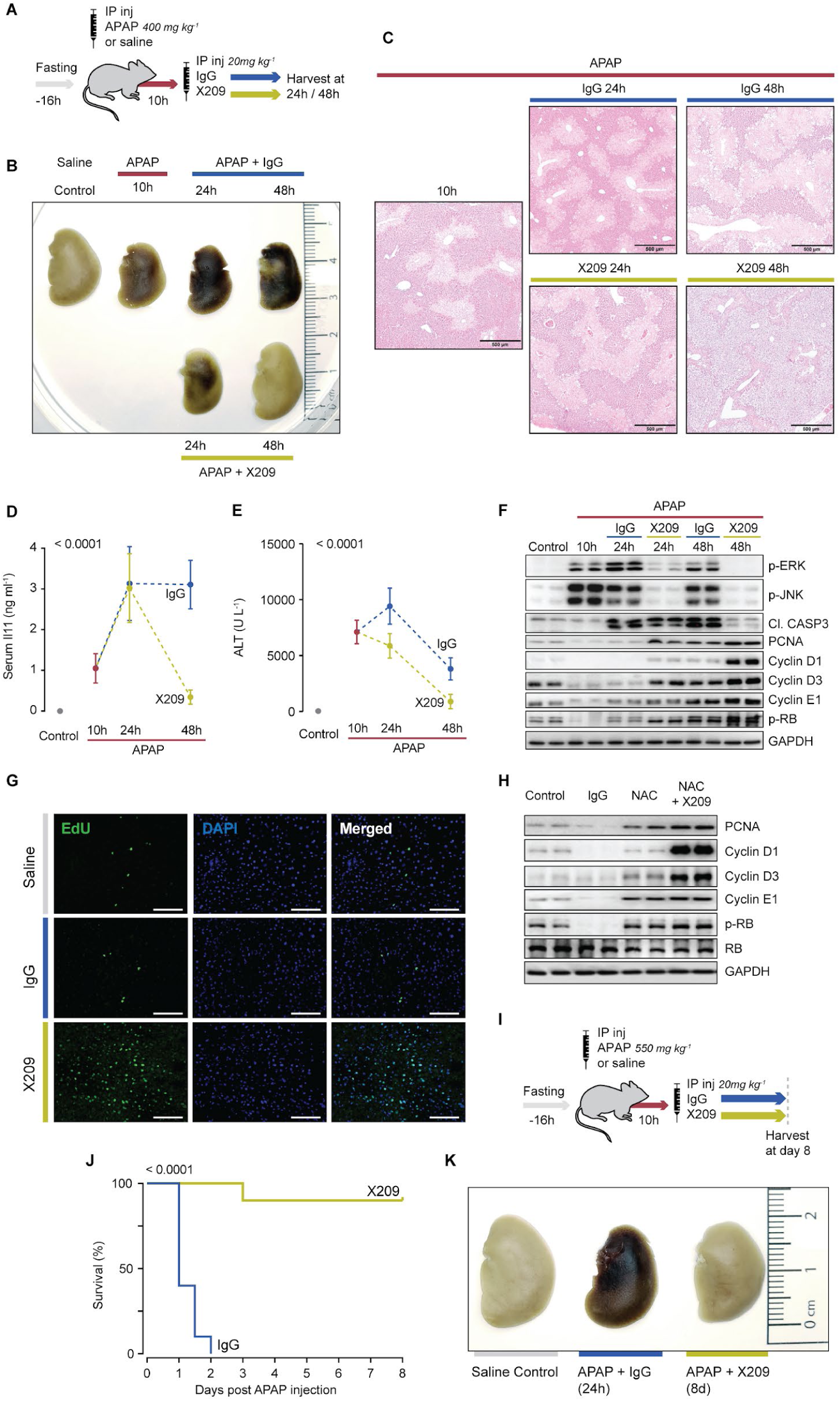
Hepatic regeneration and reversal of liver failure with late anti-IL11RA therapy. (**A**) Schematic showing late therapeutic dosing of APAP-injured mice. Overnight fasted mice were administered IgG/X209 (20 mg kg^-1^) 10h post-APAP. (**B**) Representative liver gross anatomy, (**C**) representative H&E-stained liver images (scale bars, 500 µm), (**D**) serum Il11 levels, (**E**) serum ALT levels, (**F**) western blots of p-ERK, p-JNK, Cl. CASP3, PCNA, Cyclin D1/D3/E1, and p-RB, (**G**) representative EdU-stained liver images (scale bars, 100 µm) from APAP mice receiving a late X209 dose (10h post APAP) as shown in Fig. 6A. (**H**) Western blots showing PCNA, Cyclin D1/D3/E1, p-RB protein expression levels in livers from APAP mice treated with either NAC or NAC+X209 (Schematic Fig. 5G). (**I**) Schematic of mice receiving X209 (20mg kg^-1^) treatment 10h following a lethal APAP OD (550 mg kg^-1^) for data shown in (**J-K**). (**J**) Survival curves of mice treated with either IgG or X209 10h post lethal APAP OD. (**K**) Gross liver anatomy of control (D8), IgG (24h) and X209-treated mice (D8). (**D, E**) Data are mean±SD; 2-way ANOVA; (**J**) Gehan-Breslow-Wilcoxon test.

Interventions promoting liver regeneration, which has very large potential, may provide a new means of treating AILI^12^. We therefore assessed the status of genes important for liver regeneration^10^. Inhibition of IL11 signaling was associated with a robust signature of regeneration with strong upregulation of PCNA, Cyclin D1/D3/E1, and phosphorylation of RB, as seen during regeneration following partial hepatectomy^10^. EdU injection and histological analyses showed very large numbers of nuclei with evidence of recent DNA synthesis in X209-treated mice as compared to controls (Fig. 6G). We reassessed the effects X209 given 3h post-APAP (Fig. 5I-L) to see if regeneration was also associated with inhibition of IL11 signaling at earlier time points. This proved to be the case, and the combination of X209 and NAC was more effective than NAC alone in increasing molecular markers of regeneration, notably for Cyclin D1 and D3 (Fig. 6H).

Finally, we administered X209 (20 mg kg^-1^) 10h after a higher and lethal acetaminophen dose (550 mg kg^-1^) at a time point when mice were moribund and livers undergoing fulminant necroinflammation (Fig. 6I). X209-treated mice recovered and had a 90% survival by the study end. In contrast, IgG-treated mice did not recover and succumbed with a 100% mortality within 48h, (Fig. 6J). On day 8 after the lethal dose of APAP, X209-treated mice appeared healthy with normal liver morphology and ALT levels were comparable to controls that had not received APAP (Fig. 6K, Extended Data Fig. 14A-B).

## Discussion

APAP OD is common with up to 50,000 individuals attending emergency departments every year in the UK, some who develop liver failure requiring transplantation^1^. Here, we describe the unexpected discovery that IL11, previously reported as protective against APAP-induced liver failure^17, 20^, liver ischemia^18, 21^, endotoxemia^22^ and inflammation^19^, is actually hepatotoxic and of central importance for liver failure following APAP OD.

The observation that endogenous IL11 is hepatotoxic is most surprising as over 30 publications have reported cytoprotective and/or anti-inflammatory effects of rhIL11 in rodent models of human disease (Supplementary Table 1,2). We discovered that rhIL11is a competitive inhibitor of mouse IL11 binding to IL11RA1, which overturns our understanding of the role of IL11 in AILI and liver disease more generally. This also implies that anti-IL11 therapies may be effective in additional diseases where rhIL11 had protective effects in mouse models such as rheumatoid arthritis^27^ and colitis^28^, among others (Supplementary Table 2). We highlight that, based on the erroneous assumption that rhIL11 effects in mice embodied beneficial IL11 gain-of-function, a number of clinical trials using rhIL11 were performed in patients (Supplementary Table 3).

Our study stimulates questions and has limitations. We show ERK is co-regulated with JNK post-APAP, yet ERK’s specific role in AILI is not known. Likewise, although hepatocyte-specific *Nox4* deletion is protective^26^, mice with global *Nox4* deletion are susceptible to AILI^29^. IL11 is critical for myofibroblasts^14–16^, which are also dependent on NOX4^30, 31^. In the current study, we show reduced hepatic JNK activation following IL11 inhibition, which is similar to that seen in livers of mice with hepatocyte-specific *Nox4* deletion and steatohepatitis, further suggestive of an IL11-NOX4 relationship^14, 26^. Whether anti-IL11 therapy stimulates regeneration in other organs is not known. These matters require further study.

We propose a refined mechanism for APAP toxicity whereby NAPQI damaged mitochondria produce ROS that stimulates IL11-dependent NOX4 upregulation and further sustained ROS production (Extended Data Fig. 15). This drives a dual pathology: killing hepatocytes via JNK and caspase activation and preventing hepatocyte regeneration, through mechanisms yet to be defined. The mouse model of AILI closely resembles human disease and we suggest that therapies targeting IL11 signaling might be trialed in patients with APAP-induced liver toxicity. Since IL11 neutralizing therapies are not dependent on altering APAP metabolism and specifically stimulate regeneration, they are effective much later than the current standard of care and might be particularly useful for patients presenting late to the emergency room.

## Acknowledgements

The authors would like to acknowledge the technical support of N.S.J.Ko, S.Lim, and B.L.George.

## Author contributions

A.A.W. and S.A.C. conceived and designed the study. A.A.W., J.D, S.V., B.K.S, W.W.L., S.G.S., J.T., M.W., and L.E.F., performed *in vitro* cell culture, cell biology and molecular biology experiments. A.A.W., J.D., B.N., J.Z., S.G.S, and J.T. performed *in vivo* studies. L.S.P. and C.L.D. performed mass spectrometry. S.G.S. and S.Y.L performed histology analysis. R.H. and A.H. performed surface plasmon resonance and competitive ELISA. S.P.C. performed computational analysis. J.W.D. provided critical reagents. A.A.W., J.D., P.M.Y., C.L.D., and S.A.C. analyzed the data. A.A.W., J.D., E.A., S.S., and S.A.C. prepared the manuscript with input from co-authors.

## Data and materials availability

All data are provided in the manuscript or in the supplementary materials.

---

**Extended Data Figure 1:**
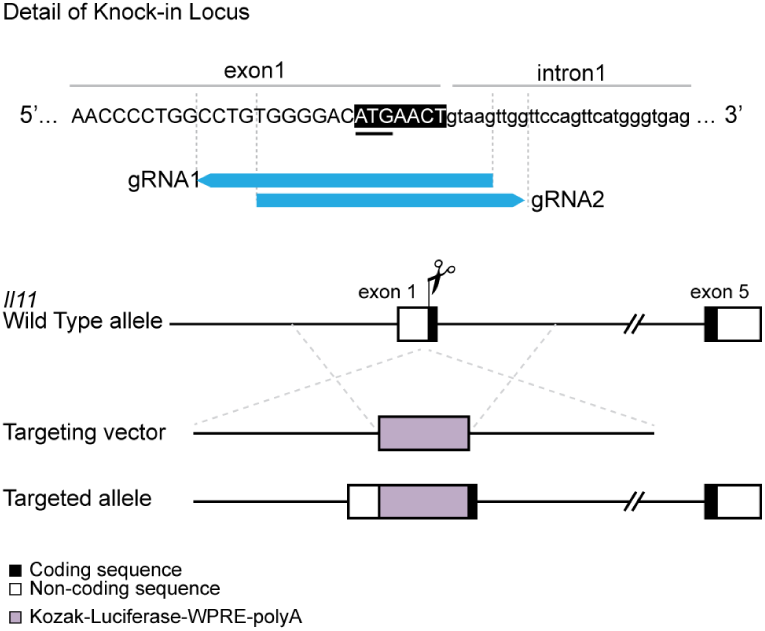
*Il11*-*Luciferase* knock-in mice. Knock-in strategy for Kozak-Luciferase-WPRE-polyA into exon 1 of *Il11* locus using CRISPR/Cas9. Woodchuck Hepatitis Virus (WHP) Posttranscriptional Regulatory Element (WPRE).

**Extended Data Figure 2:**
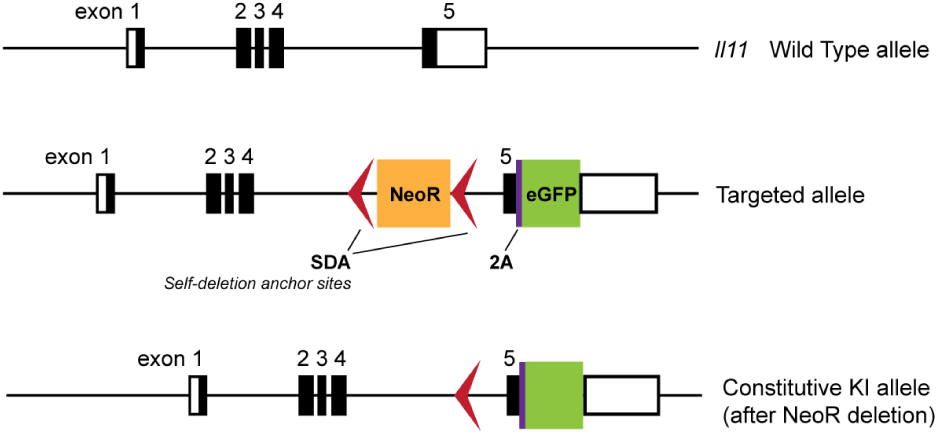
*Il11*-*EGFP* knockin mice. Knock-in strategy for *2A-EGFP* cassette into exon 5 of *Il11* gene, replacing the TGA stop codon resulting in the translation of Il11-2A-EGFP protein. The 2A linker is cleaved resulting in retention of EGFP in cells that express and secrete IL11.

**Extended Data Figure 3:**
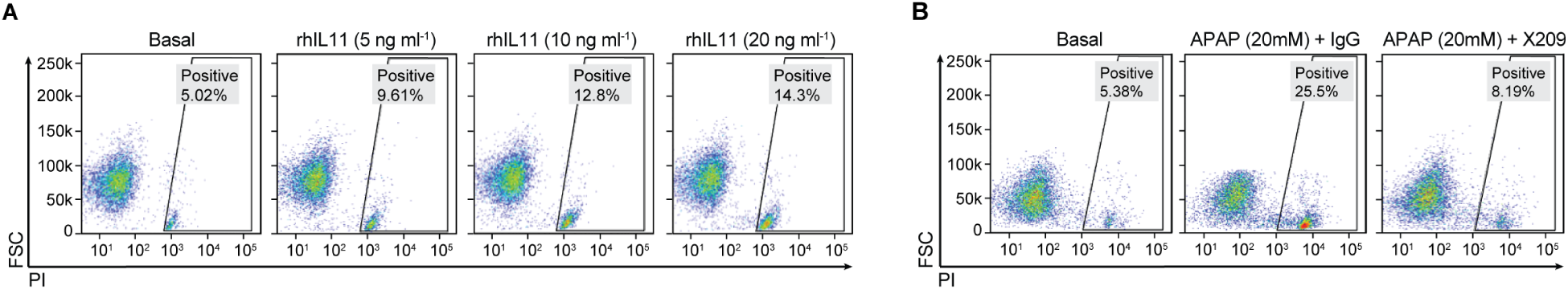
Hepatotoxic effects of IL11. Representative flow cytometry forward scatter (FSC) plots of Propidium Iodide (PI) staining of primary human hepatocytes stimulated with (**A**) increasing dose of rhIL11 and (**B**) APAP in the presence of either IgG or X209 (2 µg ml^-1^).

**Extended Data Figure 4:**
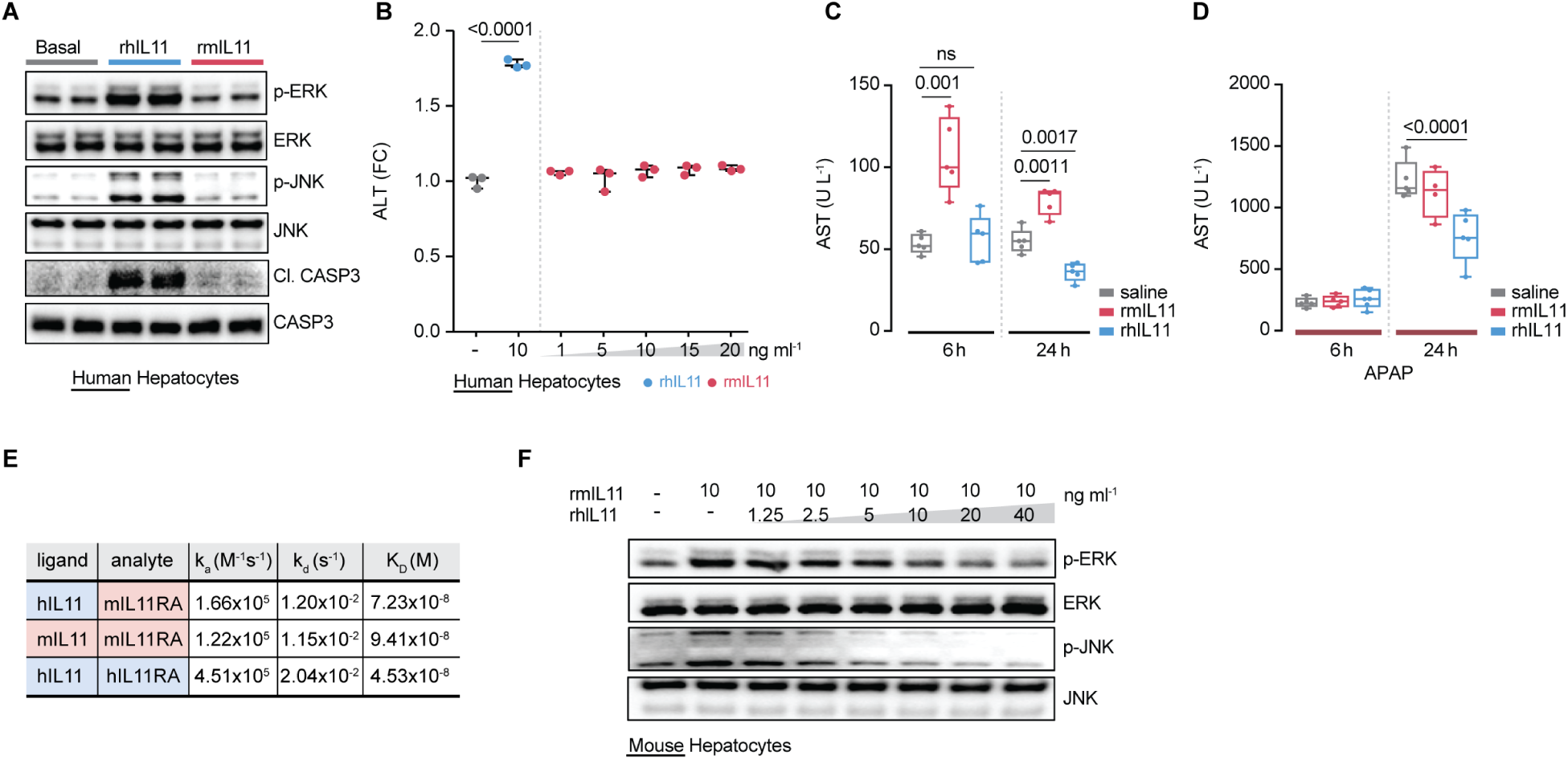
Species-specific effects of human or mouse IL11 on human or mouse hepatocytes. (**A**) Effect of recombinant human IL11 (rhIL11, 10 ng ml^-1^) or recombinant mouse IL11 (rmIL11, 10 ng ml^-1^) on ERK, JNK and CASP3 activation status in human hepatocytes. (**B**) ALT levels in the supernatant of human hepatocytes stimulated with either rhIL11 or increasing dose of rmIL11. (**C**) Effect of rhIL11 and rmIL11 treatment alone (Schematic Fig. 2C) or (**D**) with APAP administration (Schematic Fig. 2F) on serum AST levels in the mice. (**E**) Binding affinity and kinetic constants for mouse IL11RA interaction with either mouse IL11 or human IL11 and for human IL11RA interaction with human IL11. (**F**) Western blots showing dose-dependent inhibition effect of rhIL11 on p-ERK, ERK, p-JNK, JNK in mouse hepatocytes stimulated with rmIL11 (10 ng ml^-1^, 24h), (**B**) Data are shown as mean±SD; (**C,D**) Data are shown as box-and-whisker with median (middle line), 25th–75th percentiles (box), and minimum-maximum values (whiskers). (**B**) Two-tailed, Tukey-corrected Student’s *t*-test; (**C**) two-tailed Student’s *t*-test; (**D**) two-tailed Dunnett’s test. FC: fold change.

**Extended Data Figure 5:**
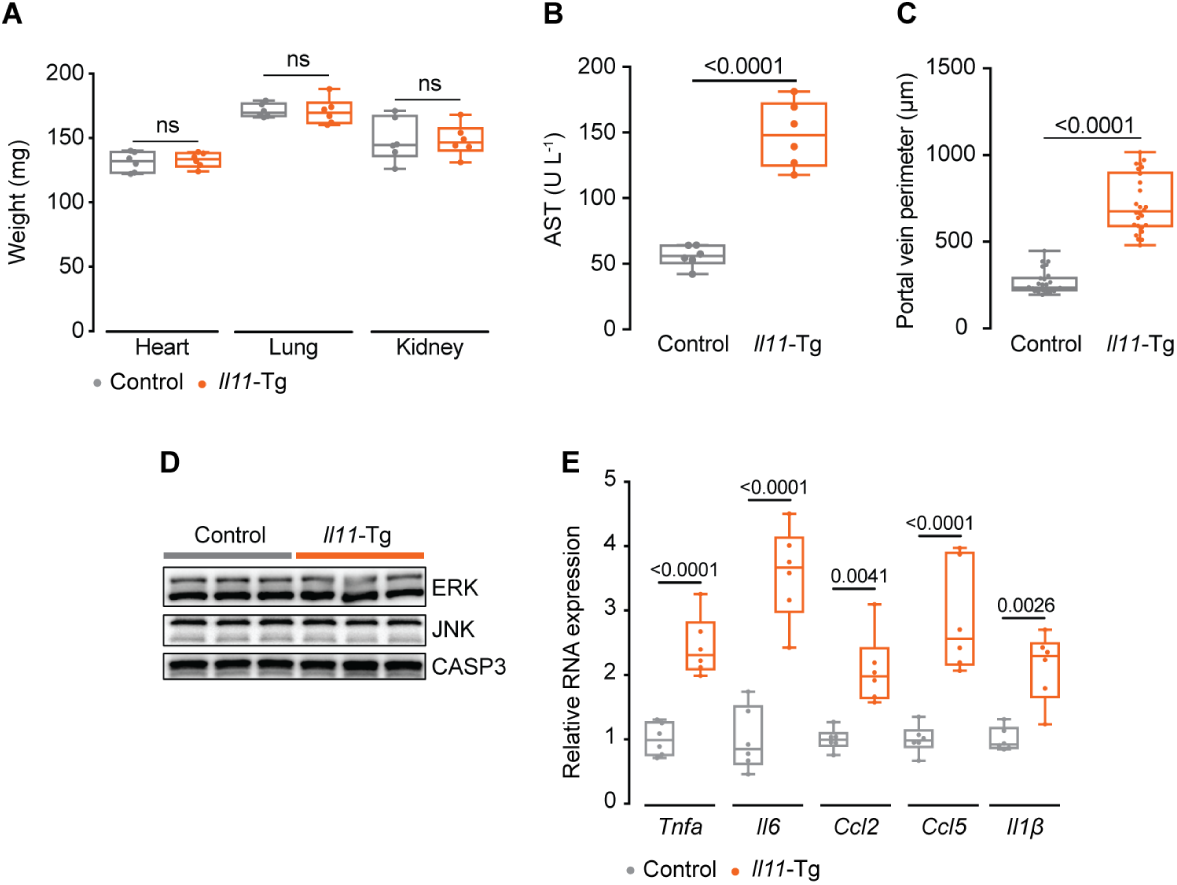
Hepatocyte-specific *Il11* overexpression causes liver necroinflammation. (**A)** Weight of heart, lung, kidney, (**B**) serum AST levels, (**C**) quantification of portal vein diameter, (**D**) Western blots of total ERK, total JNK, and CASP3, and (**E**) relative liver mRNA expression levels of pro-inflammatory markers of control and *Il11*-Tg mice (Schematic Fig. 3A). (**A-C, E**) Data are shown as box-and-whisker with median (middle line), 25th–75th percentiles (box), and minimum-maximum values (whiskers); two-tailed Student’s *t*-test.

**Extended Data Figure 6:**
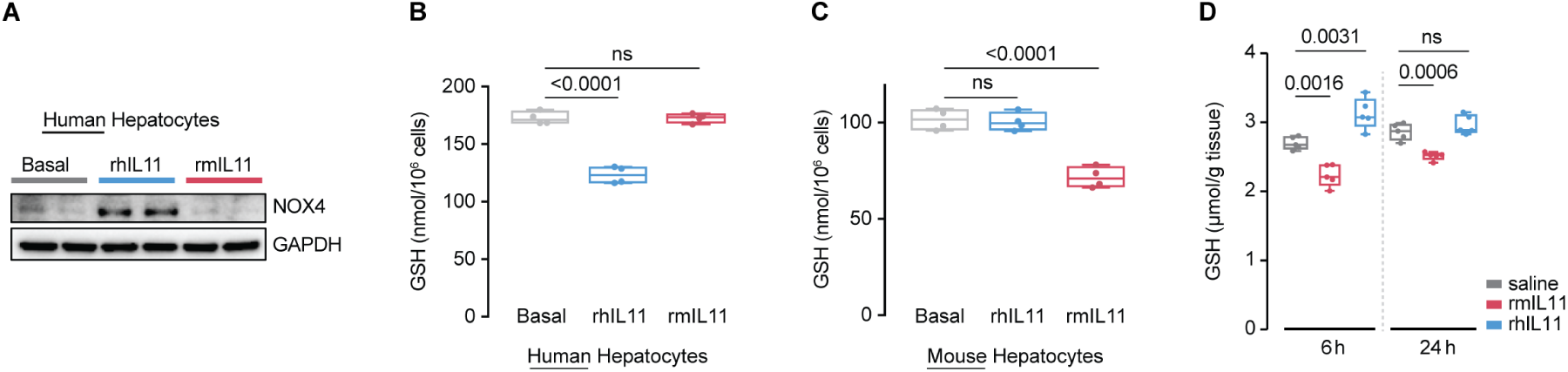
Only species-specific IL11 induces NOX4 and glutathione depletion in hepatocytes. Effect of rhIL11 and rmIL11 (10 ng ml^-1^) on (**A**) NOX4 protein expression, (**B**) GSH levels in human hepatocytes, (**C**) GSH levels in mouse hepatocytes. (**D**) Hepatic GSH levels following rhIL11 or rmIL11 administration to mice (Schematic Fig. 2C) (**B-D**) Data are shown as box-and-whisker with median (middle line), 25th–75th percentiles (box), and minimum-maximum values (whiskers); two-tailed Dunnett’s test.

**Extended Data Figure 7:**
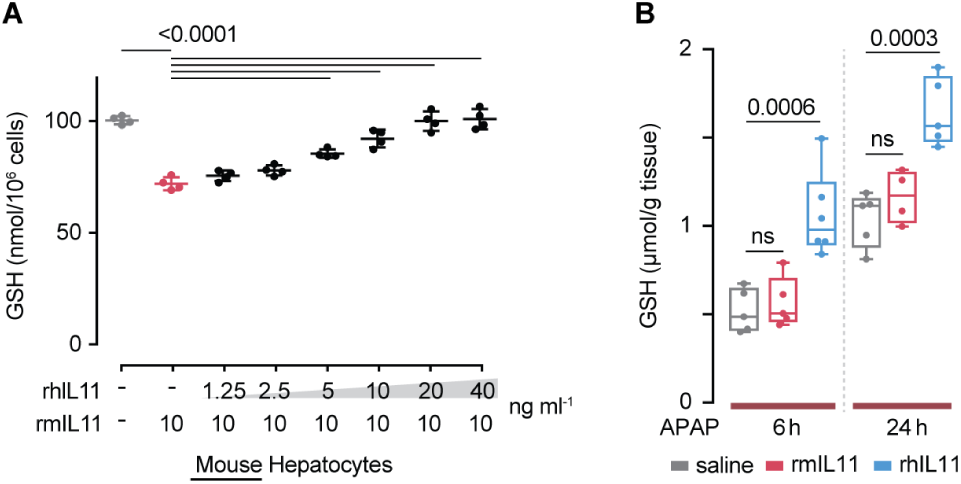
Recombinant human IL11 (rhIL11) restores GSH levels in injured mouse liver. (**A**) Dose-dependent inhibition effect of rhIL11 on GSH levels in primary mouse hepatocytes stimulated with rmIL11; two-tailed, Tukey-corrected Student’s *t*-test. (**B**) Effect of rhIL11 or rmIL11 on murine hepatic GSH levels following APAP injury, as shown in Schematic Fig. 2F; two-tailed Dunnett’s test. (**A, B**) Data are shown as box-and-whisker with median (middle line), 25th–75th percentiles (box), and minimum-maximum values (whiskers).

**Extended Data Figure 8:**
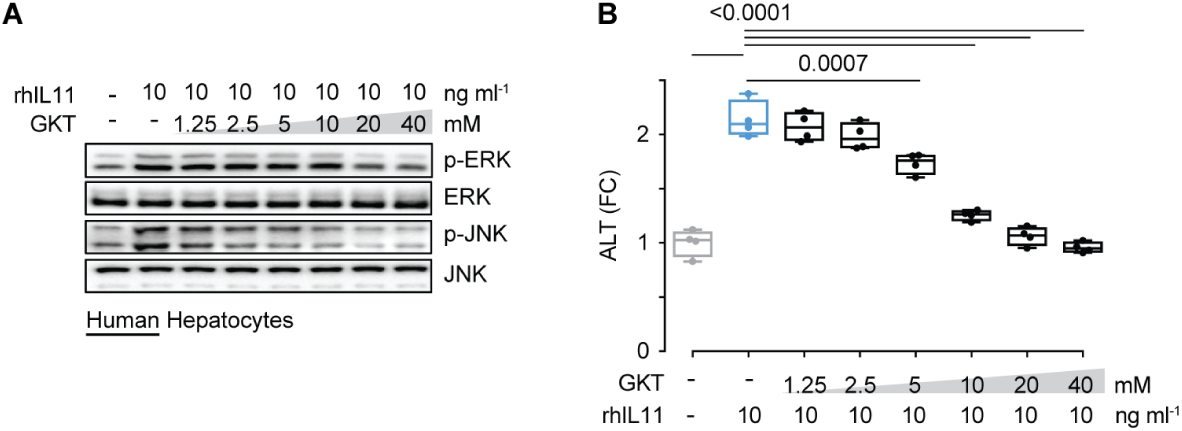
The NOX4 inhibitor GKT-137831 prevents the hepatotoxic effects of IL11. Dose-dependent inhibition effect of GKT-137831, a NOX4 inhibitor, on (**A**) ERK and JNK activation and on (**B**) ALT secretion from human hepatocytes stimulated with rhIL11 (10 ng ml^-1^, 24h). (**B**) Data are shown as box-and-whisker with median (middle line), 25th–75th percentiles (box), and minimum-maximum values (whiskers); two-tailed, Tukey-corrected Student’s *t*-test. FC: fold change.

**Extended Data Figure 9:**
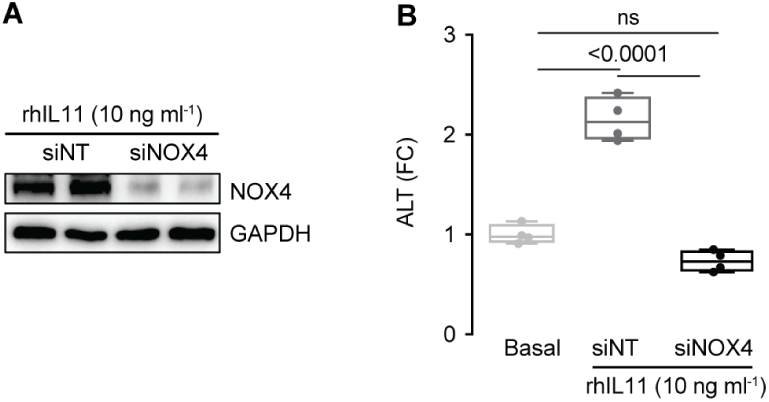
NOX4 is critical for the hepatotoxic effect of IL11. (**A**) Western blots showing the knockdown efficiency of siNOX4. (**B**) Effect of siNOX4 on rhIL11-induced primary human hepatocyte death and release of ALT. (**A-B**) rhIL11 (10 ng ml^-1^), siNT (non-targeting siRNA control)/siNOX4 (50 nM); 24h; data are shown as box-and-whisker with median (middle line), 25th–75th percentiles (box), and minimum-maximum values (whiskers); two-tailed, Tukey-corrected Student’s *t*-test. FC: fold change.

**Extended Data Figure 10:**
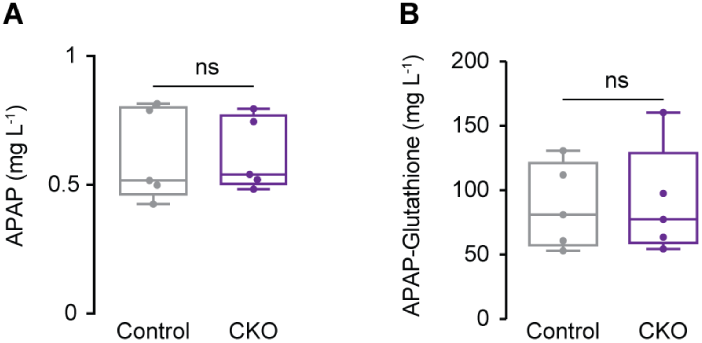
Control and CKO mice have similar serum levels of APAP and APAP-Glutathione 24h after APAP administration. LC-MS/MS Quantification of (**A**) APAP and (**B**) APAP-Glutathione in the serum of control and CKO mice. Data are shown as box-and-whisker with median (middle line), 25th–75th percentiles (box), and minimum-maximum values (whiskers); two-tailed Student’s *t*-test

**Extended Data Figure 11:**
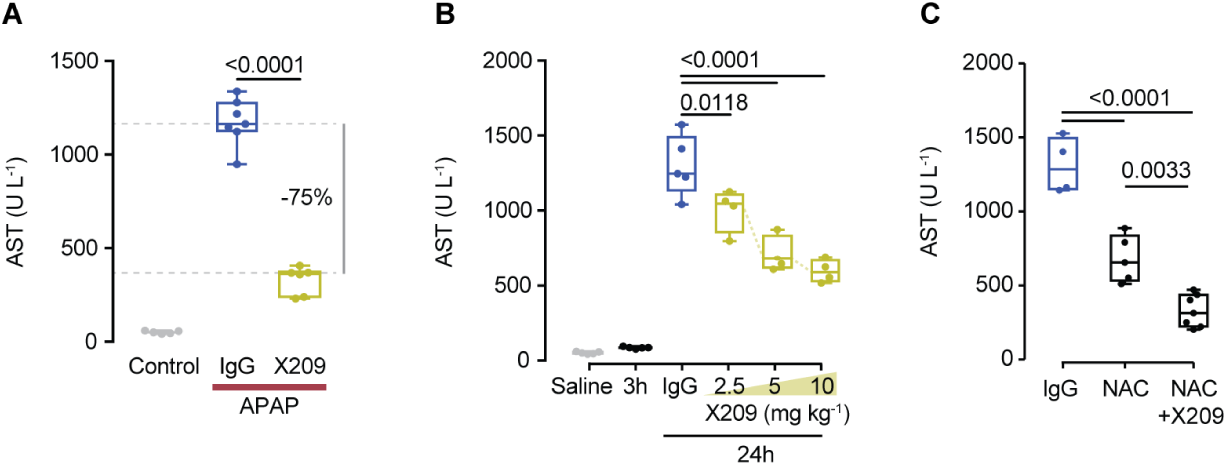
Anti-IL11RA antibody (X209) lowers serum AST after APAP injury. (**A**) Serum AST levels in saline and APAP mice receiving a preventive dose of X209 (10 mg kg^-1^), 16h prior to APAP (Schematic Fig. 5A). (**B**) Dose-dependent effect of X209 on serum AST levels in APAP mice receiving a therapeutic dose of X209, 3h post APAP administration (Schematic Fig. 5D, the values of saline are the same as those used in **S11A**). (**C**) Serum AST levels in mice treated with NAC (500 mg kg^-1^) alone or in combination with X209 (5 mg kg^-1^) 3h after APAP injury (Schematic Fig. 5G). (**A-C**) Data are shown as box-and-whisker with median (middle line), 25th–75th percentiles (box), and minimum-maximum values (whiskers); two-tailed, Tukey-corrected Student’s *t*-test.

**Extended Data Figure 12:**
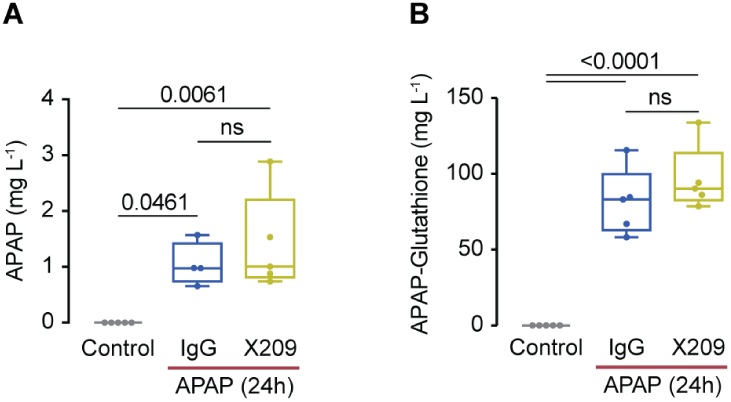
Serum levels of APAP and APAP-Glutathione in the mice serum 24h post APAP OD. LC-MS/MS Quantification of (**A**) APAP and (**B**) APAP-Glutathione in saline control mice, and in IgG and X209-treated mice 24h following APAP administration. Data are shown as box-and-whisker with median (middle line), 25th–75th percentiles (box), and minimum-maximum values (whiskers); Two-tailed, Tukey-corrected Student’s *t*-test.

**Extended Data Figure 13:**
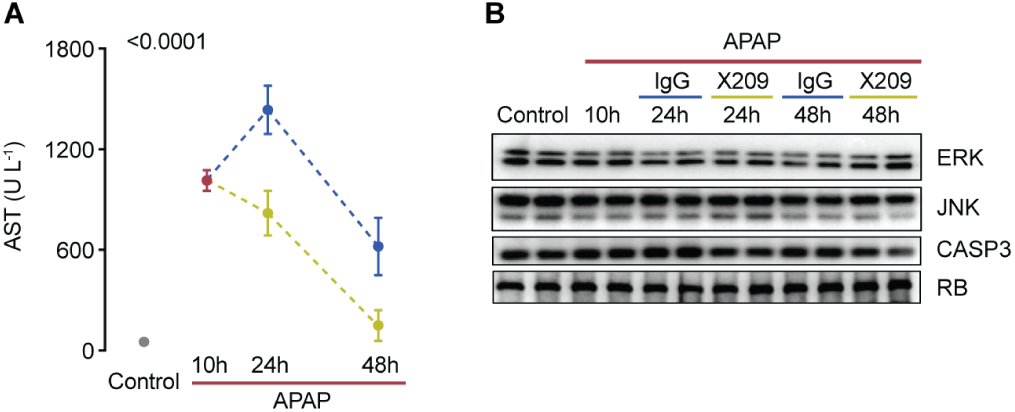
X209 reverses APAP-induced liver damage. (**A**) Serum AST levels and (**B**) Western blots showing hepatic content of total ERK, JNK, CASP3, and RB from mice in reversal experimental groups as shown in schematic Fig. 6A.

**Extended Data Figure 14:**
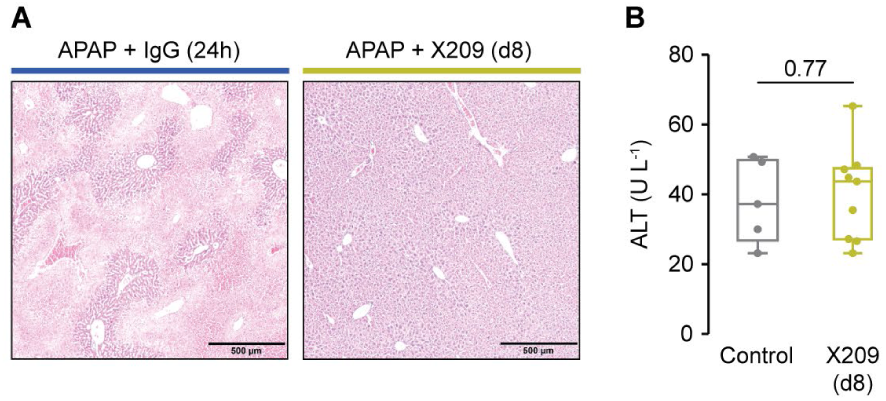
Recovery of X209-treated mice following administration of lethal APAP dose. (**A**) Representative H&E images (scale bars, 500 µm) of livers from IgG (24h post APAP) and X209-treated mice (D8 post APAP). (**B**) Serum ALT levels of saline-control and X209-treated mice (D8 post APAP).

**Extended Data Figure 15:**
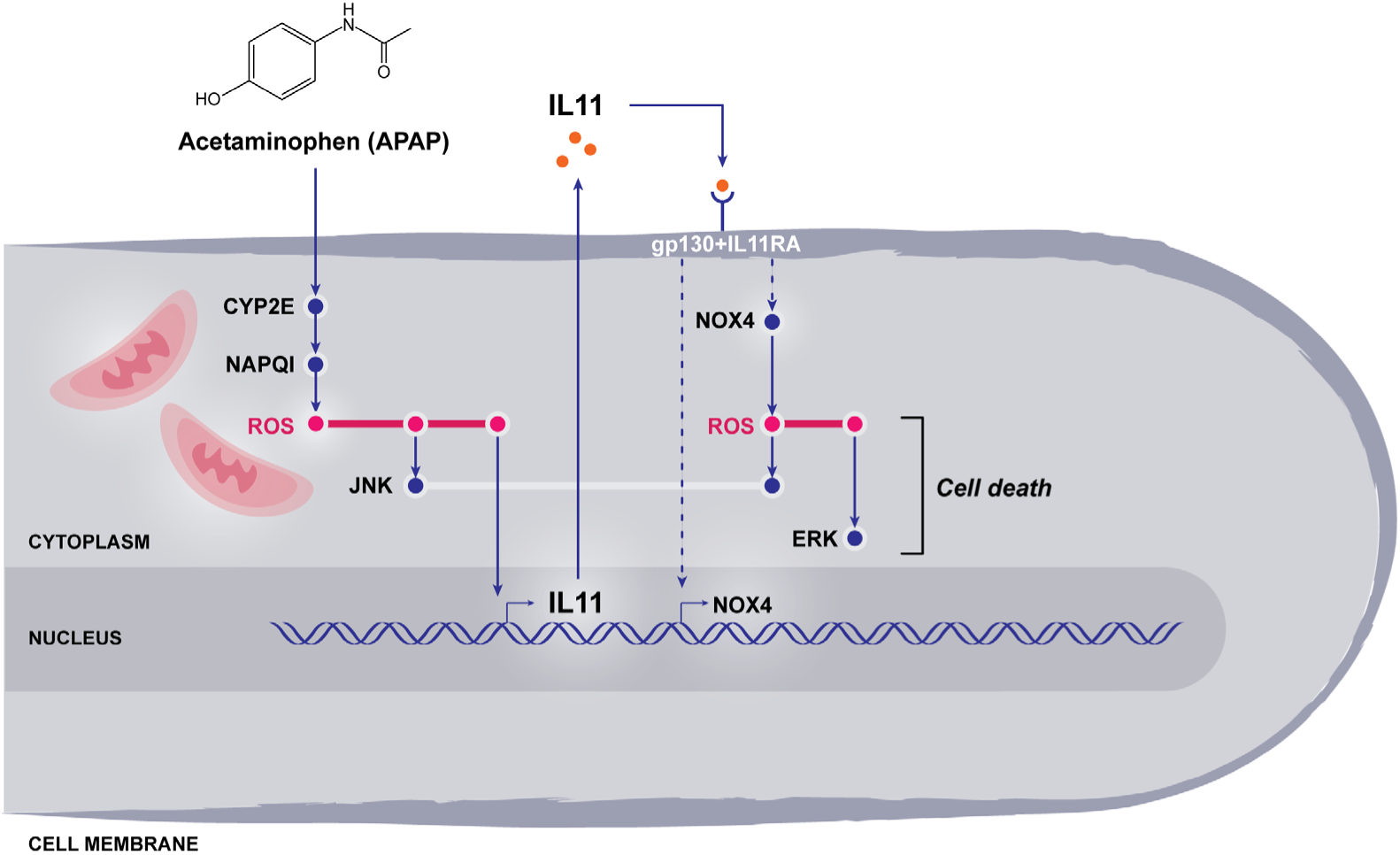
Proposed mechanism and role of IL11 in APAP-induced hepatotoxicity. Metabolizing APAP in the liver leads to ROS production via NAPQI and triggers IL11 secretion. The autocrine IL11 signaling loops on hepatocytes and continues to generate ROS via NOX4, which drives sustained cell death and limits hepatic regeneration independently of APAP and its metabolites. If the IL11 pathway is blocked either genetically or therapeutically, hepatocyte cell death can be prevented and liver regeneration is restored.

**Table 1.**
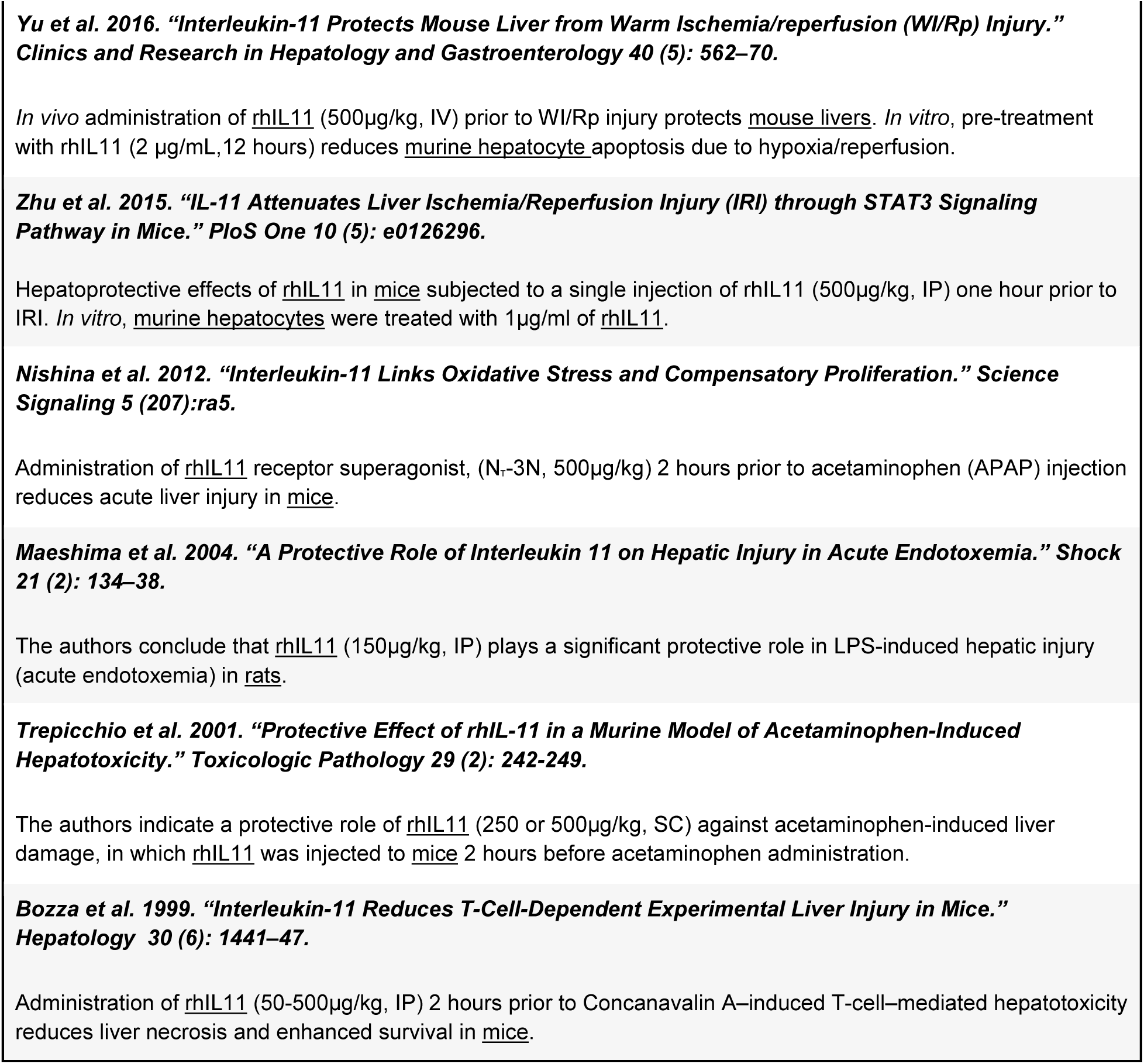
List of publications showing protective effects of recombinant human IL11 (rhIL11) in rodent models of liver injury

**Supplementary Table 2.** List of publications showing protective and/or anti-inflammatory effects of rhIL11 in other rodent disease models

|  |  |
| --- | --- |
| **Bowel** | **Gibson et al. 2010. "Interleukin-11 Reduces TLR4-Induced Colitis in TLR2-Deficient Mice and Restores Intestinal STAT3 Signaling." *Gastroenterology* 139 (4): 1277–88.**<br>The authors report that administration of rhIL11 (5µg/kg, IP) ameliorates infection colitis and is cytoprotective in TLR2-deficient mice. |
|  | **Boerma et al. 2007. "Local Administration of Interleukin-11 Ameliorates Intestinal Radiation Injury in Rats." *Cancer Research* 67 (19): 9501–6.**<br>The authors conclude that IL11 ameliorates early intestinal radiation injury, in which rats were given daily injections of rhIL11 (2 mg/kg/d) from 2 days prior until 2 weeks after irradiation. |
|  | **Opal et al. 2003. "Orally Administered Recombinant Human Interleukin-11 Is Protective in Experimental Neutropenic Sepsis." *The Journal of Infectious Diseases* 187 (1): 70–76.**<br>The authors suggest that IL11 maintains epithelial cell integrity during cytoreductive chemotherapy by cyclophosphamide based on effects observed in rats receiving daily oral administration of rhIL11 (0.5mg/kg/day), starting from 1 day before the first dose of cyclophosphamide for a total of 12 days. |
|  | **Ropeleski et al. 2003. "Interleukin-11-Induced Heat Shock Protein 25 Confers Intestinal Epithelial-Specific Cytoprotection from Oxidant Stress." *Gastroenterology* 124 (5): 1358–68.**<br>The authors conclude that IL11 confers epithelial-specific cytoprotection during intestinal epithelial injury. Rat, mouse and canine cell lines (IEC-18, YAMC, NIH3T3, MDCK-HR) were stimulated with high (50-100ng/ml) levels of rhIL11. |
|  | **Greenwood-Van Meerveld et al 2000. "Recombinant Human Interleukin-11 Modulates Ion Transport and Mucosal Inflammation in the Small Intestine and Colon." *Laboratory Investigation; a Journal of Technical Methods and Pathology* 80 (8): 1269–80.**<br>The authors conclude that during intestinal inflammation IL11 acts as a modulator of epithelial transport or as an anti-inflammatory cytokine based on effects of rhIL11 on rat mucosal sheets (10-10,000 ng/ml) and in rats (33µg/kg, alternate days for 1 or 2 weeks). |
|  | **Du et al 1997. "Protective Effects of Interleukin-11 in a Murine Model of Ischemic Bowel Necrosis." *American Journal of Physiology-Gastrointestinal and Liver Physiology*.**<br>Administration of rhIL11 (250 µg/kg/day) for 3 days prior to and for 7 days post bowel ischemia induction confers a protective effect against ischemic bowel necrosis and the authors suggest its use as a treatment for gastrointestinal mucosal diseases. |
|  | **Orazi et al. 1996. "Interleukin-11 Prevents Apoptosis and Accelerates Recovery of Small Intestinal Mucosa in Mice Treated with Combined Chemotherapy and Radiation." *Laboratory Investigation; a Journal of Technical Methods and Pathology* 75 (1): 33–42.**<br>Administration of rhIL11 (250 µg/kg) promotes recovery from chemotherapy and radiation-induced damage to the mice small intestinal mucosa. |
|  | **Potten et al 1996. "Protection of the Small Intestinal Clonogenic Stem Cells from Radiation-Induced Damage by Pretreatment with Interleukin 11 Also Increases Murine Survival Time." *Stem Cells*.1996 14(4):452-9.**<br>RhIL11 (100µg/kg, SC), administered to mice prior to and after cytotoxic exposure, protects clonogenic cells in intestinal crypts and increases murine survival times following radiation exposure. |
|  | **Qiu et al. 1996. "Protection by Recombinant Human Interleukin-11 against Experimental TNB-Induced Colitis in Rats." *Digestive Diseases and Sciences* 41 (8): 1625–30.**<br>The authors describe protective effects of rhIL11 in trinitrobenzene sulfonic acid-induced colitis in rats. Rats were injected daily with rhIL11 (100, 300, or 1000µg/kg, SC) 3 days before, or daily for 3-7-14 days after TNB administration. |
|  | **Du et al. 1994. "A Bone Marrow Stromal-Derived Growth Factor, Interleukin-11, Stimulates Recovery of Small Intestinal Mucosal Cells after Cytoablative Therapy." *Blood* 83 (1): 33–37.**<br>Administration of rhIL11 (250µg/kg/d, SC) promotes recovery of small intestinal mucosa following combination radiation and chemotherapy in mice. |
| Heart | **Tamura et al. 2018. "The Cardioprotective Effect of Interleukin-11 against Ischemia-Reperfusion Injury in a Heart Donor Model." *Annals of Cardiothoracic Surgery* 7 (1):99-105.**<br>Administration of rhIL11 (18µg/ml, IV, 10 minutes prior to heart collection) preserves heart function and lower apoptosis index in rat following ex vivo model of cold ischemia. |
|  | **Obana et al. 2012. "Therapeutic Administration of IL-11 Exhibits the Postconditioning Effects against Ischemia-Reperfusion Injury via STAT3 in the Heart." *American Journal of Physiology. Heart and Circulatory Physiology* 303 (5): H569–77.**<br>Administration of rhIL11 (20 µg/kg, IV at the start of reperfusion) prevents adverse cardiac remodeling and apoptosis after ischemia reperfusion injury-induced acute myocardial infarction in mice |
|  | **Obana et al. 2010. "Therapeutic Activation of Signal Transducer and Activator of Transcription 3 by Interleukin-11 Ameliorates Cardiac Fibrosis after Myocardial Infarction." *Circulation* 121 (5): 684–91.**<br>Administration of rhIL11 (8µg/kg, IV) 24 hours following left coronary artery ligation-induced myocardial infarction (MI) and then consecutively every 24 hours for 4 days reduces post-MI scar volume in mice. |
|  | **Kimura et al. 2007. "Identification of Cardiac Myocytes as the Target of Interleukin 11, a Cardioprotective Cytokine." *Cytokine* 38 (2):107-115**<br>The authors conclude that IL11 is a cardioprotective based on effects of rhIL11 (8µg/kg) administered to mouse 15 hours prior to cardiac ischemia-reperfusion |
| Immune System | **Bozza et al. 2001. "Interleukin-11 Modulates Th1/Th2 Cytokine Production from Activated CD4 T Cells." *Journal of Interferon & Cytokine Research* 21(1):21-30.**<br>The authors state that IL11 acts directly on activated murine CD4+ve T-cells and modulates, not represses, the immune response following stimulation with rhIL11 (1-500 ng/ml). |
|  | **Opal et al. 2000. "Recombinant Human Interleukin-11 Has Anti-inflammatory Actions Yet Does Not Exacerbate Systemic Listeria Infection." *The Journal of Infectious Diseases* 181(2): 754-756** |
|  | Daily administration of rhIL11 (150 mg/kg, IV) for 7 days prior to Listeria infection reduces interferon-γ levels. Interestingly, the authors stated that inflammatory markers IL-6/IFN-γ trend down after anti-IL11mAb (10mg/kg) treatment. |
|  | **Hill et al. 1998. "Interleukin-11 Promotes T Cell Polarization and Prevents Acute Graft-versus-Host Disease after Allogeneic Bone Marrow Transplantation." *The Journal of Clinical Investigation* 102 (1): 115–23.**<br>The authors conclude that IL11 prevents Graft-vs-Host-Disease (GVHD) via T Cell polarization, based on experiments in which a high dose of rhIL11 (250μg/kg, SC, twice daily) was injected into a murine model of GVHD. |
|  | **Sonis et al. 1997. "Mitigating Effects of Interleukin 11 on Consecutive Courses of 5-Fluorouracil-Induced Ulcerative Mucositis in Hamsters." *Cytokine* 9 (8): 605–12.**<br>Administration of rhIL11 (50-100μg/animal/day, SC) protects from 5-fluorouracil-induced ulcerative mucositis in hamsters. |
|  | **Trepicchio et al. 1997. "IL-11 Regulates Macrophage Effector Function through the Inhibition of Nuclear Factor-kappaB." *Journal of Immunology* 159 (11): 5661–70.**<br>The authors conclude that IL11 inhibits the secretion of pro-inflammatory cytokines by macrophages; murine primary macrophages were treated with rhIL11 (10-100 ng/ml). |
|  | **Trepicchio et al. 1996. "Recombinant Human IL-11 Attenuates the Inflammatory Response through down-Regulation of Proinflammatory Cytokine Release and Nitric Oxide Production." *Journal of Immunology* 157 (8): 3627–34.**<br>The authors report that IL11 reduces levels of TNF-α, IL-1β and IFN-γ in the serum of LPS-treated mice and in LPS-stimulated macrophage media. Mice and murine macrophages were treated with rhIL11 (500μg/kg or 10-100 ng/ml, respectively). |
| Joint | **Anguita et al. 1999. "Selective Anti-Inflammatory Action of Interleukin-11 in Murine Lyme Disease: Arthritis Decreases While Carditis Persists." *The Journal of Infectious Diseases* 179 (3): 734–37.**<br>Administration of rhIL11(0.1-2μg/mouse/day, 5 days/week for 3 weeks) reduces arthritis, but not carditis, in *Borrelia burgdorferi*-infected mice (a murine model of Lyme disease). |
|  | **Walmsley et al. 1998. "An Anti-Inflammatory Role for Interleukin-11 in Established Murine Collagen-Induced Arthritis." *Immunology* 95 (1): 31–37.**<br>Daily administration of rhIL11 (0.3-100μg/mouse/day, IP, 10 days) reduces inflammation in a murine model of collagen-induced arthritis. |
| Kidney | **Lee et al. 2012. "Interleukin-11 Protects against Renal Ischemia and Reperfusion Injury." *American Journal of Physiology. Renal Physiology* 303 (8): F1216–24.**<br>The authors conclude that IL11 is renoprotective based on pre-treatment (10 minutes prior to IR) and post-treatment (30-60 minutes following IR) effects of rhIL11 and PEGylated rhIL11 (100-1000μg/kg, IP) in mice. |
|  | **Stangou et al. 2011. "Effect of IL-11 on Glomerular Expression of TGF-Beta and Extracellular Matrix in Nephrotoxic Nephritis in Wistar Kyoto Rats." *Journal of Nephrology* 24 (1): 106–111.** |
|  | Administration of rhIL11 (800-1360µg/kg, IP) 2 hours prior to nephrotoxic nephritis and then once daily for 6 days suppresses ECM deposition in rats. |
| Lung | **Sheridan et al 1999. "Interleukin-11 Attenuates Pulmonary Inflammation and Vasomotor Dysfunction in Endotoxin-Induced Lung Injury." *The American Journal of Physiology* 277 (5): L861–67.**<br>The authors conclude that rhIL11 (200mg/kg, IP) exerts an anti-inflammatory activity that protects against LPS-induced lung injury and lethality in rats |
|  | **Waxman et al. 1998. "Targeted Lung Expression of Interleukin-11 Enhances Murine Tolerance of 100% Oxygen and Diminishes Hyperoxia-Induced DNA Fragmentation." *J. Clin. Invest.* 101(9):1970-1982**<br>The authors conclude that IL11 protects from hyperoxic-induced lung injury, based on the effects of lung-specific human IL11 overexpression in mice. |

**Supplementary Table 3.**
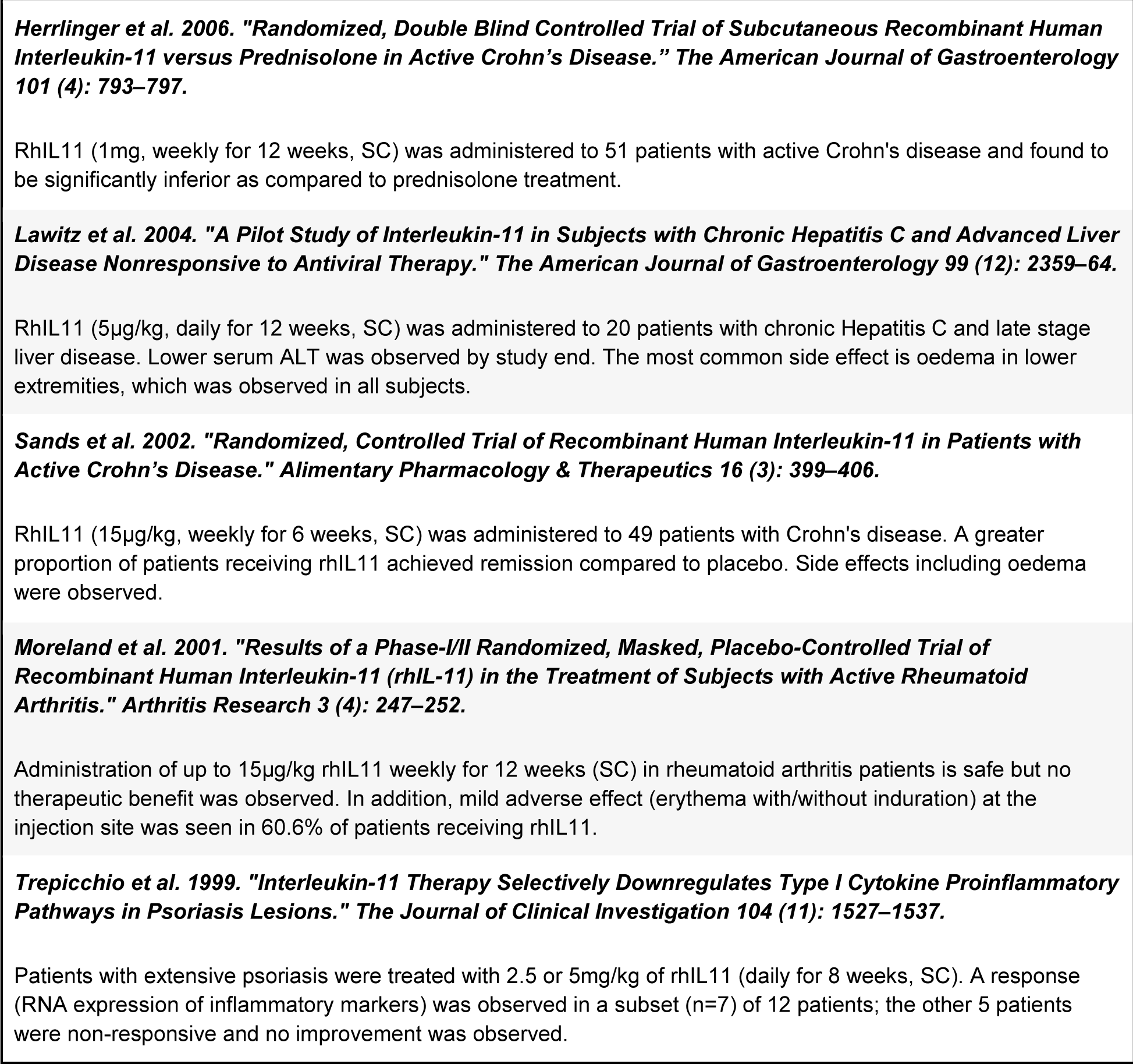
List of publications from clinical trials where rhIL11 was administered to patients, based mainly on an inferred protective effect of rhIL11 use in rodent models of disease.

## References and notes

1. Bernal, W. & Wendon, J. Acute liver failure. The New England journal of medicine 370, 1170–1171 (2014).

2. Lee, W. M. et al. Intravenous N-acetylcysteine improves transplant-free survival in early stage non-acetaminophen acute liver failure. Gastroenterology 137, 856–64, 864.e1 (2009).

3. Jaeschke, H. Acetaminophen: Dose-Dependent Drug Hepatotoxicity and Acute Liver Failure in Patients. Dig. Dis. 33, 464–471 (2015).

4. Chiew, A. L., Gluud, C., Brok, J. & Buckley, N. A. Interventions for paracetamol (acetaminophen) overdose. Cochrane Database Syst. Rev. 2, CD003328 (2018).

5. Win, S. et al. New insights into the role and mechanism of c-Jun-N-terminal kinase signaling in the pathobiology of liver diseases. Hepatology 67, 2013–2024 (2018).

6. Zhang, H. et al. Reduction of liver Fas expression by an antisense oligonucleotide protects mice from fulminant hepatitis. Nat. Biotechnol. 18, 862–867 (2000).

7. Schwabe, R. F. & Luedde, T. Apoptosis and necroptosis in the liver: a matter of life and death. Nat. Rev. Gastroenterol. Hepatol. 15, 738–752 (2018).

8. Gunawan, B. K. et al.c-Jun N-Terminal Kinase Plays a Major Role in Murine Acetaminophen Hepatotoxicity. Gastroenterology 131, 165–178 (2006).

9. Xie, Y. et al. Inhibitor of apoptosis signal-regulating kinase 1 protects against acetaminophen-induced liver injury. Toxicol. Appl. Pharmacol. 286, 1–9 (2015).

10. Sekiya, S. & Suzuki, A. Glycogen synthase kinase 3 β-dependent Snail degradation directs hepatocyte proliferation in normal liver regeneration. Proc. Natl. Acad. Sci. U. S. A. 108, 11175–11180 (2011).

11. Marcos, A. et al. Liver regeneration and function in donor and recipient after right lobe adult to adult living donor liver transplantation. Transplantation 69, 1375–1379 (2000).

12. Bhushan, B. & Apte, U. Liver Regeneration after Acetaminophen Hepatotoxicity: Mechanisms and Therapeutic Opportunities. Am. J. Pathol. 189, 719–729 (2019).

13. Michalopoulos, G. K. Hepatostat: Liver regeneration and normal liver tissue maintenance. Hepatology 65, 1384–1392 (2017).

14. Widjaja, A. A. et al. Inhibiting Interleukin 11 Signaling Reduces Hepatocyte Death and Liver Fibrosis, Inflammation, and Steatosis in Mouse Models of Non-Alcoholic Steatohepatitis. Gastroenterology (2019). doi:10.1053/j.gastro.2019.05.002

15. Schafer, S. et al. IL-11 is a crucial determinant of cardiovascular fibrosis. Nature 552, 110–115 (2017).

16. Cook, S. et al. IL-11 is a therapeutic target in idiopathic pulmonary fibrosis. (2018). doi:10.1101/336537

17. Nishina, T. et al. Interleukin-11 links oxidative stress and compensatory proliferation. Sci. Signal. 5, ra5 (2012).

18. Zhu, M. et al. IL-11 Attenuates Liver Ischemia/Reperfusion Injury (IRI) through STAT3 Signaling Pathway in Mice. PLoS One 10, e0126296 (2015).

19. Bozza, M. et al. Interleukin-11 reduces T-cell-dependent experimental liver injury in mice. Hepatology 30, 1441–1447 (1999).

20. Trepicchio, W. L., Bozza, M., Bouchard, P. & Dorner, A. J. Protective effect of rhIL-11 in a murine model of acetaminophen-induced hepatotoxicity. Toxicol. Pathol. 29, 242–249 (2001).

21. Yu, J., Feng, Z., Tan, L., Pu, L. & Kong, L. Interleukin-11 protects mouse liver from warm ischemia/reperfusion (WI/Rp) injury. Clin. Res. Hepatol. Gastroenterol. 40, 562–570 (2016).

22. Maeshima, K. et al. A protective role of interleukin 11 on hepatic injury in acute endotoxemia. Shock 21, 134–138 (2004).

23. Mühl, H. STAT3, a key parameter of cytokine-driven tissue protection during sterile inflammation--the case of experimental acetaminophen (Paracetamol)-induced liver damage. Front. Immunol. 7, 163 (2016).

24. Schleinkofer, K. et al. Identification of the Domain in the Human Interleukin-11Receptorthat Mediates Ligand Binding. available online at http://www.idealibrary.com on J. Mol. Biol. 306, 263–274 (2001).

25. Denton, C. P. et al. Therapeutic interleukin-6 blockade reverses transforming growth factor-beta pathway activation in dermal fibroblasts: insights from the faSScinate clinical trial in systemic sclerosis. Ann. Rheum. Dis. 77, 1362–1371 (2018).

26. Bettaieb, A. et al. Hepatocyte Nicotinamide Adenine Dinucleotide Phosphate Reduced Oxidase 4 Regulates Stress Signaling, Fibrosis, and Insulin Sensitivity During Development of Steatohepatitis in Mice. Gastroenterology 149, 468–80.e10 (2015).

27. Walmsley, M., Butler, D. M., Marinova-Mutafchieva, L. & Feldmann, M. An anti-inflammatory role for interleukin-11 in established murine collagen-induced arthritis. Immunology 95, 31–37 (1998).

28. Qiu, B. S., Pfeiffer, C. J. & Keith, J. C. Protection by recombinant human interleukin-11 against experimental TNB-induced colitis in rats. Digestive Diseases and Sciences 41, 1625–1630 (1996).

29. Murray, T. V. A. et al. NADPH oxidase 4 regulates homocysteine metabolism and protects against acetaminophen-induced liver damage in mice. Free Radic. Biol. Med. 89, 918–930 (2015).

30. Hecker, L. et al. NADPH oxidase-4 mediates myofibroblast activation and fibrogenic responses to lung injury. Nat. Med. 15, 1077–1081 (2009).

31. Wermuth, P. J., Mendoza, F. A. & Jimenez, S. A. Abrogation of transforming growth factor-β-induced tissue fibrosis in mice with a global genetic deletion of Nox4. Lab. Invest. 99, 470–482 (2019).

