## Supplementary material for "Redefining Interleukin 11 as a regeneration-limiting hepatotoxin": Methods

**Materials and methods**

**Antibodies**

Cleaved Caspase 3 (9664, CST), Caspase 3 (9662, CST), Cyclin D1 (55506, CST), Cyclin D3 (2936, CST), Cyclin E1 (20808, CST), p-ERK1/2 (4370, CST), ERK1/2 (4695, CST), GAPDH (2118, CST), GFP (ab6673, Abcam), IgG (Aldevron), p-JNK (4668, CST), JNK (9258, CST), neutralizing anti-IL11RA (X209, Aldevron; *in vivo* study), IL11RA (130920, Santa Cruz; WB), NOX4 (110-58849, Novus Biologicals), PCNA (13110, CST), p-RB (8516, CST), RB (9313, CST), anti-rabbit HRP (7074, CST), anti-mouse HRP (7076, CST), anti-rabbit Alexa Fluor 488 (ab150077, Abcam), anti-rabbit Alexa Fluor 647 (ab150079, Abcam), anti-mouse Alexa Fluor 488 (ab150113, Abcam), anti-goat Alexa Fluor 488 (ab150129, Abcam) .

**Recombinant proteins**

Recombinant human IL11 (rhIL11, UniProtKB:P20809, Genscript), recombinant mouse IL11 (rmIL11, UniProtKB: P47873, Genscript), human IL11RA (10252-H08H, SinoBiological), mouse IL11RA (50075-M08H, SinoBiological).

**Chemical**

Acetaminophen (APAP, A3035, Sigma), DAPI (D1306, ThermoFisher Scientifics), D-Luciferin (L6882, Sigma), GKT-137831 (17764, Cayman Chemical), N-Acetyl-L-Csyteine (NAC, A7250, Sigma).

**Reagents for LC-MS/MS**

Reference Standard acetaminophen (APAP, P0300000, Sigma), internal standard (IS) acetaminophen-d4 (APAP-D4, A161222, Toronto Research Chemicals), IS acetaminophen glutathione (APAP GLUT, A161223, Toronto Research Chemicals), Acetronitrile (900667, Sigma), Ammonium formate (A115-50, Sigma), Formic acid (F0507, Sigma), mouse serum (IGMSCD1SER50ML, i-DNA Biotechnology). All chemicals, reagents and solvents were of LC–MS grade quality.

**Animal models**

Animal procedures were approved and conducted in accordance with the SingHealth Institutional Animal Care and Use Committee (IACUC). All mice were provided food and water *ad libitum*, except in the fasting period, during which only water was provided *ad libitum*.

***Mouse models of Acetaminophen poisoning***

Prior to APAP, 12-14 weeks old male mice (C57BL6/NTAC, unless otherwise specified in the main text or figure legends) were fasted overnight. Mice were then given a severe (400 mg $kg^{-1}$) or lethal (550 mg $kg^{-1}$) dose of APAP by intraperitoneal (IP) administration. Mice were administered anti-IL11RA (X209) or IgG isotype control antibody at different times and doses, depending on the experiments as outlined in the main text or figure legends. Mice were euthanized at various time points post APAP, from 10h to 8 days, as outlined in the main text or figure legends.

***Il11-Luciferase mice***

The mouse *Il11* gene consists of 5 exons, with the ATG start codon in exon 1 and TGA stop codon in exon 5. Three transcripts of mouse *Il11* have been identified (ENSMUSG00000004371): transcript *Il11-201* is the longest and encodes a 199aa pro-peptide, whereas transcripts *Il11-202* and *Il11-203* contain an alternative first exon, and are both predicted to encode a shorter 140 aa isoform that lacks the signal peptide. Using the

CRISPR/Cas9 technique, a Kozak-Luciferase-WPRE-polyA sequence was introduced to replace the ATG start codon in exon 1 of *Il11-201* (ENSMUST00000094892.11), resulting in translational disruption of this specific transcript. Single guide RNAs (sgRNAs) with recognition sites in exon 1 along with Cas9 and the targeting construct containing a Kozak-Luciferase-WPRE-polyA sequence were microinjected into fertilized zygotes and subsequently transferred into pseudopregnant mice (Shanghai Model Organisms Center, Inc). Insertion of the luciferase cassette into the *Il11* gene locus was verified by sequencing. Mutant *Il11-Luciferase* offsprings were generated on a C57BL/6 background and identified by genotyping to detect the insertion of the luciferase construct in exon 1, using primers which amplify a 818 bp region corresponding to the wildtype *Il11* allele (5'-GGAGGGAGGGGACGCCAATGACC-3' and 5'- TCTGCCTCCCCTGCCTGTTTCTCG -3'), and a second set of primers that amplifies a 928 bp region corresponding to the targeted allele containing the luciferase construct (5'-AATTCCGTGGTGTTGTCG-3' and 5'-TCTGCCTCCCCTGCCTGTTTCTCG-3'). Heterozygous *Il11-Luciferase* were subjected to APAP-induced liver injury as described above. After 24h, mice were injected intraperitoneally with 150 mg $kg^{-1}$ of D-Luciferin in PBS and bioluminescence images of the liver were subsequently acquired using the IVIS Lumina System (Perkin Elmer), according to the manufacturer's instructions.

***Il11-EGFP mice***

Transgenic mice with *EGFP* constitutively knocked-in to the *Il11* gene were generated by Cyagen Biosciences Inc. Briefly, knockin mice were generated to contain a *2A-EGFP* cassette inserted into exon 5, which replaces the TGA stop codon sequence, and translation of the targeted transcript would give rise to full-length IL11 pro-peptide and EGFP separated by a 2A self cleaving peptide linker. The targeting vector homology arms of *Il11* gene, containing a Neo cassette inserted into intron 4 (flanked by SDA: self-deletion anchor sites) and a *2A-EGFP* cassette inserted into exon 5, were generated by PCR using BAC clones from the C57BL/6 library. C57BL/6 ES cells were used for gene targeting and successfully targeted clones were injected into C57BL/6 albino embryos, which were then re-implanted into CD-1 pseudo-pregnant females. Founder animals were identified by their coat color and germline transmission was confirmed by breeding with C57BL/6 females and subsequent genotyping of the offsprings. Genotyping primers were designed to amplify selected regions of intron 4, spanning the Neo cassette SDA sites, according to the following primer sequences:

(5'- GAAATGAGAGCCTAGAGTCCAGAG-3' and
5'- GAGGCTTGGAAGAATGCACAATTA-3').

***Hepatocyte-specific Il11 overexpressing mice (Il11-*Tg*)***

We previously generated mice in which mouse *Il11* cDNA was introduced into the *Rosa26* locus under the control of *loxP-Stop-loxP* sites to allow for cell-type specific overexpression of Il11 following Cre recombinase-mediated excision[1]. These animals are made available at The Jackson Laboratory (*C57BL/6N-Gt(ROSA)26Sortm1(CAG-Il11)Cook/J*). To induce the specific expression of *Il11* in hepatocytes, heterozygous *Il11-Rosa26* mice were injected intravenously (IV) with either $4x10^{11}$ genome copies in PBS/mouse (VectorBiolabs) of AAV8-ALB-Null (Control) AAV8-ALB-Cre (*Il11*-Tg). Livers and serum were assessed after three weeks.

***Hepatocyte-specific Il11ra1 deleted mice***

We recently generated and validated *Il11ra1*-floxed mice, in which exons 4 to 7 of the *Il11ra1* gene were flanked by loxP sites, allowing for the spatial and temporal deletion of *Il11ra1* upon Cre recombinase-mediated excision[2]. To induce the specific deletion of *Il11ra1* in hepatocytes, homozygous *Il11ra1*-floxed mice were IV injected with AAV8-ALB-Cre virus ($4x10^{11}$ genome copies in PBS/mouse, VectorBiolabs) via the tail vein. A similar amount of AAV8-ALB-Null virus were injected into homozygous *Il11ra1*-floxed mice as controls. The AAV8 treated mice were allowed to recover for three weeks prior to APAP injury. Knockdown efficiency was determined by Western blotting of hepatic IL11RA.

**<u>Cell culture</u>**

Both primary human and mouse hepatocytes were grown and maintained at 37°C and 5% $CO_2$. The growth medium was renewed every 2–3 days and cells were passaged at 80% confluence, using standard trypsinization techniques. All experiments were carried out at low cell passage (P1-P3). Stimulated cells were compared to unstimulated cells that have been grown for the same duration under the same conditions, but without the stimuli.

***Primary human hepatocytes***

Human hepatocytes (5200, ScienCell) were maintained in hepatocyte medium (520, ScienCell) supplemented with 2% fetal bovine serum, 1% Penicillin-streptomycin. Cells were serum-starved for 16h prior to respective stimulations, as outlined in the main text or figure legends, that were performed in serum-free hepatocyte media for 24h.

***Primary mouse hepatocytes***

Mouse hepatocytes (ABC-TC3928, AcceGen Biotech) were maintained in mouse hepatocyte medium (ABC-TM3928, AcceGen Biotech) supplemented with 1% Penicillin-streptomycin. Cells were stimulated with different treatment conditions, as outlined in the main text or figure legends for 24h.

**<u>siRNA transfection</u>**

Primary human hepatocytes were seeded at 60–70% confluency in 6-well plate, 16h before transfection. Cells were transfected with 50 nM of *NOX4* siRNA (ON-TARGETplus SMARTpool siRNA, L-010194-00-0005, Dharmacon) or control siRNA (D-001810-10-05, Dharmacon) for 24h at 37°C in OptiMEM (31985070, Thermo Fisher) containing Lipofectamine RNAiMAX Transfection Reagent (13778150, Thermo Fisher). Transfected cells were then stimulated with rhIL11 for 24h. Knockdown efficiency was determined by immunoblotting of NOX4.

**<u>Flow cytometry</u>**

Primary human hepatocytes ($5\times10^5$) were stained using FITC Annexin V/Dead Cell Apoptosis Kit (V13242, Thermo Fisher), according to the manufacturer's instructions. $PI^{+ve}$ cells were quantified with the flow cytometer (Fortessa, BD Biosciences) and analyzed with FlowJo version 7 software (TreeStar).

**<u>Colorimetric assays</u>**

The levels of alanine transaminase (ALT) or aspartate aminotransferase (AST) in mouse serum and hepatocyte supernatant were measured using ALT Activity (ab105134, Abcam) or AST (ab105135, Abcam) Assay Kits. Liver glutathione sulfhydryl (GSH) measurements were performed using Glutathione Colorimetric Detection Kit (EIAGSHC, Thermo Fisher). All colorimetric assays were performed according to the manufacturer's protocol.

**<u>Enzyme-linked immunosorbent assay (ELISA)</u>**

The levels of IL11 in mouse serum and hepatocyte supernatant were quantified using Mouse IL-11 DuoSet (DY418 and DY008, R&D Systems) and Human IL11 Quantikine ELISA kit D1100, R&D Systems), respectively, according to the manufacturer's protocol.

***Competitive ELISA***

Mouse IL11RA (1 µg $ml^{-1}$ in PBS) was coated on a 96-well plate (overnight at 4°C) and then blocked with blocking buffer (1%BSA in PBS containing 0.05% Tween20). Biotinylated mouse Il11 was prepared using Lightning-Link Rapid Biotin type A kit (Expedeon) according to the manufacturer's instructions. RhIL11 or rmIl11 was two-fold serially diluted in blocking buffer (starting at 5 μg $ml^{-1}$) and mixed with 0.01 μg $ml^{-1}$ biotinylated mouse IL11. The mixture of biotinylated mouse IL 11 and either rhIL11 or rmIL11 was added into the coated plate and incubated for 1h at RT. Color development was performed by adding Streptavidin-HRP (1:1000 in blocking buffer) and TMB chromogen solution (002023, ThermoFisher Scientific).

**Immunoblotting**

Western blots were carried out from hepatocyte and liver tissue lysates. Hepatocytes and tissues were homogenized in radioimmunoprecipitation assay (RIPA) buffer containing protease and phosphatase inhibitors (Thermo Fisher), followed by centrifugation to clear the lysate. Protein concentrations were determined by Bradford assay (Bio-Rad). Equal amounts of protein lysates were separated by SDS-PAGE, transferred to PVDF membrane, and subjected to immunoblot analysis for the indicated primary antibodies. Proteins were visualized using the ECL detection system (Pierce) with the appropriate secondary antibodies.

**Quantitative polymerase chain reaction (qPCR)**

Total RNA was extracted from either the snap-frozen liver tissues or hepatocyte lysates using Trizol (Invitrogen) followed by RNeasy column (Qiagen) purification. cDNAs were synthesized with iScript™ cDNA synthesis kit (Bio-Rad) according to manufacturer's instructions. Gene expression analysis was performed on duplicate samples with either TaqMan (Applied Biosystems) or fast SYBR green (Qiagen) technology using StepOnePlus™ (Applied Biosystem) over 40 cycles. Expression data were normalized to *GAPDH* mRNA expression and fold change was calculated using $2^{-\Delta\Delta Ct}$ method. The sequences of specific TaqMan probes and SYBR green primers are available upon request.

**Surface plasmon resonance (SPR)**

SPR measurements were performed on a BIAcore T200 (GE Healthcare) at 25ºC. Buffers were degassed and filter-sterilized through 0.2 μm filters prior to use. RhIL11 or rmIl11 was immobilized onto a carboxymethylated dextran (CM5) sensor chip using standard amine coupling chemistry. For kinetic analysis, a concentration series (3.125 nM to 100 nM) of human IL11RA or mouse Il11ra was injected over the rhIL11, rmIl11 and reference surfaces at a flow rate of 40 μl $min^{-1}$. All the analytes were dissolved in HBS-EP+ (BR100669, GE Healthcare) containing 1 mg $ml^{-1}$ BSA. The association and dissociation were measured for 150s and 200s respectively. After each analyte injection, the surface was regenerated by a 30s injection of Glycine pH2.5, followed by a 5 min stabilisation period. All sensorgrams were aligned and double-referenced. Affinity and kinetic constants were determined by fitting the corrected sensorgrams with the 1:1 Langmuir model using BIAevaluation v3.0 software (GE Healthcare). The equilibrium binding constant $K_D$ was determined by the ratio of the binding rate constants $k_d/k_a$.

**Histology**

***Hematoxylin&Eosin (H&E) staining***

Livers were fixed for 48h at room temperature (RT) in 10% neutral-buffered formalin (NBF), dehydrated, embedded in paraffin blocks and sectioned at 7 μm. Sections were stained with H&E according to standard protocol and examined by light microscopy.

***EdU staining***

Livers were rinsed in cold PBS and patted dry with a lint free paper and cryo-molded in OCT compound (4583, Tissue-Tek®). After the OCT compound is frozen, liver specimens were wrapped in aluminium foil and stored in -80°C. Cryo-embedded livers were cryosectioned (-20°C) at 7 μm thickness and allowed to dry on the slides for 1h before proceeding to EdU detection using Baseclick's EdU IV Imaging Kit 488L (BCK488-IV-IM-L) according to the manufacturer's protocol.

***Immunofluorescence staining***

Livers were processed and frozen as mentioned above (EdU staining section). Frozen liver tissues were sectioned at 7 μm at -20°C and left to dry for 1h (RT). Liver sections were fixed in cold acetone for 15 min prior to brief PBS washes, permeabilized with 0.1% TritonX-100 (T8787, Sigma), and blocked with 2.5% normal goat serum (S-1012, Vector Labs) for 1h (RT). Liver sections were incubated with GFP (1:500) and Caspase 3 (1:1000) primary antibodies overnight (4°C), followed by incubation with the appropriate Alexa Fluor 488/647 secondary antibodies (1:250) for 1h (RT). DAPI was used to stain the nuclei prior to imaging by fluorescence microscope (Leica).

**LC-MS/MS**

Mouse serum samples (20 μL), calibrators and QCs were transferred into a deep well 96-well plate, then spiked with 50 μL of 10 μg $l^{-1}$ of APAP-D4 heavy isotope standards. After treating with 360 μL of ice-cold Acetonitrile containing 0.1% Formic acid, the plate was mixed (1000 rpm $min^{-1}$, 10 min), followed by centrifugation (2270g, 50 min, 4°C). 140 μL of the supernatant was carefully transferred to a 96-microwell plate and loaded into the auto-sampler for analysis by LC-MS/MS. Ion counts were then normalized against that of the heavy isotope standard, before using the standard curve for quantification. Liquid chromatographic (LC) separation of the biomarkers was carried out on an Agilent 1290 Infinity II LC system (Agilent Technologies) with PEEK coated SeQuant®ZIC®-cHILIC 3mm,100Å 100 x 2.1 mm HPLC column (Merck Pte Ltd) maintained at 40°C. The organic solvent was Acetonitrile containing 0.1% Formic acid (Solvent A) and the aqueous solvent was 20 mM Ammonium Formate pH 4.0 (Solvent B). A linear LC gradient on Binary Pump A (G7120A, Agilent Technologies) was set up with percentage of Solvent B as follows: 10% at 0 min, 70% at 9 min, 70% at 11 min, and 10% between 11.1 and 11.5min, with a flow rate of 0.4 ml $min^{-1}$. The column was further equilibrated for 11.5 min with 10% Solvent B. An additional high speed pump, Binary Pump B, together with a Quick-Change valve head, 2-position/10-port, 1,300 bar (5067-4240, Agilent), were utilized to reduce the cycle times by automated alternating column regeneration. Percentage of Solvent B on BinaryPump B was maintained at 10% with a flow rate of 0.3 ml $min^{-1}$. For mass detection, the LC eluent is connected to an Agilent 6495 Triple Quadrupole MS system (G6495A, Agilent Technologies) operated with the electrospray source in either positive or negative ionization mode. The electrospray ionization source conditions were as follows: capillary voltage of 4.0 kV, nozzle voltage of 500 V, iFunnel parameter high/low pressure RF of 90 V, nebulizer pressure of 60 psi, gas temperature of 290°C, sheath gas temperature of 350°C, Nebulizer was 35 psi, and sheath gas flow of 12 l $min^{-1}$. The multiple reaction monitoring (MRM) conditions used for APAP and APAP-D4 were 152.1→110 with Collision Energy (CE) of 16eV, Collision Accelerator Voltage (CAV) of 5 V and 156→114 with CE of 8eV and CAV of 5 V, respectively. The MRM used for APAP-Glutathione was 457.1→140 with Collision Energy (CE) of 42eV, Collision Accelerator Voltage (CAV) of 5 V.

***Calibration and linearity***

Nine-point calibration curves were obtained by fortifying drug-free mouse serum with working solutions of APAP and APAP-Glutathione. The final concentrations of APAP were 0.32, 0.46, 2.6, 5.2, 10.3, 20.6, 41.3, 82.5 and 330 mg $l^{-1}$ (low QC: 1.29 mg $l^{-1}$; high QC: 165

mg $l^{-1}$). The final concentrations of APAP-Glutathione were 0.244, 0.49, 0.98, 1.95, 3.91, 7.81, 15.6, 62.5, 125 and 250 mg $l^{-1}$ (low QC: 1.95 mg $l^{-1}$; high QC of 31.3 mg $l^{-1}$). Standard curves corresponded to peak area ratios of each analyte to IS using weighted linear least-squares regression (1/x2) for APAP and (1/x) for APAP-Glutathione, the linearity coefficients of determination (r2) were 0.97807145 and 0.99655914, respectively. The precision and accuracy of the assay in the mice serum samples were determined as described previously[3].

**Statistical analysis**

Statistical analyses were performed using GraphPad Prism software (version 6.07). P values were corrected for multiple testing according to Dunnett's (when several experimental groups were compared to one condition), Tukey (when several conditions were compared to each other within one experiment), Sidak (when several conditions from 2 different genotypes were compared to each other). Analysis for two parameters for comparison of two different groups were performed by two-way ANOVA. Survival curves were analyzed by Gehan-Breslow-Wilcoxon test. The criterion for statistical significance was $P < 0.05$.

### References


1. Schafer, S. *et al.* IL-11 is a crucial determinant of cardiovascular fibrosis. *Nature* **552**, 110–115 (2017).
2. Ng, B. *et al.* Fibroblast-specific IL11 signaling is required for lung fibrosis and inflammation. *bioRxiv* 801852 (2019). doi:10.1101/801852
3. Gicquel, T., Aubert, J., Lepage, S., Fromenty, B. & Morel, I. Quantitative Analysis of Acetaminophen and its Primary Metabolites in Small Plasma Volumes by Liquid Chromatography-Tandem Mass Spectrometry. *Journal of Analytical Toxicology* **37**, 110–116 (2013).
